# Neuronal overexpression of Nrf2 reduces dystrophic neurites in 5XFAD Alzheimer’s disease model mice

**DOI:** 10.64898/2026.03.16.711596

**Authors:** Katherine R. Sadleir, Karen P. Gomez, Sidhanth Chandra, Makenna L. Ley, Ammaarah W. Khatri, Joanna Guo, Yunlu Xue, Constance L. Cepko, Robert Vassar

## Abstract

**Background:** The hallmark lesions of the Alzheimer’s disease (AD) brain are amyloid plaques consisting of the β-amyloid protein and neurofibrillary tangles comprised of hyperphosphorylated, aggregated tau protein, which both cause neuronal dysfunction and loss. One goal of neuroprotective therapies is to maintain normal neuronal function and survival in the presence of toxic pathologies such as plaques and tangles. A potential neuroprotective target is nuclear factor erythroid 2-related factor 2 (Nrf2) transcription factor, which regulates the expression of many antioxidant and detoxification genes. Nrf2 mRNA is decreased in AD brains, and deletion of the Nrf2 gene causes increased BACE1 and Aβ production and worsened cognitive deficits in amyloid pathology mouse models. Overexpression of Nrf2 in astrocytes has been shown to be protective against neurodegeneration, but the role of Nrf2 is neurons is unclear.

**Methods:** We overexpressed Nrf2 from birth in neurons of 5XFAD amyloid pathology model mice using AAV8, hypothesizing that neuronal Nrf2 overexpression decreases cortical neuron loss and reduces plaque load by decreasing BACE1 levels. We quantified protein levels by immunoblot and neuropathology by immunofluorescent staining, using two-way ANOVA to measure differences between genotypes and AAV treatments. To assess genetic changes, we performed bulk mRNA seq.

**Results:** While neuronal overexpression of Nrf2 in 5XFAD mice did not prevent neuronal loss as measured by NeuN labeling, decrease neuroinflammation by Iba1 or GFAP labeling, or reduce amyloid load by Aβ antibody or methoxy-XO4 staining, we show that increased Nrf2 expression reduces BACE1 protein levels, especially in swollen axonal dystrophic neurites around amyloid plaques. Other proteins that accumulate in dystrophic neurites were also reduced, indicating decreased dystrophic neurites overall. Immunoblot analysis suggested increased autophagy was unlikely to play a role, while bulk mRNA sequencing indicated changes in lipid metabolism and microtubule stability may have contributed to reduced dystrophic neurite formation.

**Conclusions:** Dystrophic neurites impair action potential conductance and contribute to tau seeding and spreading. Their reduction by neuronal Nrf2 overexpression may protect neurons against these pathologic changes. Further study of the mechanisms by which Nrf2 reduces dystrophic neurites may lead to therapeutic strategies that can limit neuritic damage caused by cerebral amyloid accumulation.

## Background

Alzheimer’s disease is characterized by progressive cognitive decline caused by the accumulation of two key pathologic lesions in the brain, extracellular amyloid plaques made of aggregated β-amyloid peptides (Aβ) and intracellular neurofibrillary tangles (NFT), consisting of hyperphosphorylated aggregates of tau protein, and are accompanied by neuroinflammation. With advances in brain imaging, such as positron emission tomography (PET) for amyloid and tau pathologies, it has become clear that amyloid plaque deposition precedes tau tangle formation by many years [1–11]. Additionally, amyloid positivity by imaging or cerebral spinal fluid measurements predicts onset of NFT accumulation and cognitive decline [3, 10, 12, 13]. The amyloid cascade hypothesis posits that toxic Aβ species (e.g., oligomers, fibrils, protofibrils) lead to tau hyperphosphorylation, fragmentation, and neurofibrillary tangle (NFT) formation (reviewed in) [14], but the mechanistic connection between Aβ and tau pathologies in AD is not well understood. One current hypothesis is that dystrophic neurites, swollen axons in contact with plaques, accumulate hyperphosphorylated, toxic forms of tau that begin to aggregate and then propagate the misfolded template to other synaptically connected cells [15–17].

While drugs targeting β-amyloid plaques and tau neurofibrillary tangles, are in development and three anti-Aβ antibodies, aducanamab, lecanamab and donanemab [18, 19] received approval from the United States FDA, therapies targeting other mechanisms of AD are desperately needed. Anti-Aβ antibodies slow but do not halt AD, benefit only early AD, and can have serious side effects. Therefore, discovering new, safe drugs that benefit all stages of AD is of paramount importance.

The cognitive decline and memory impairment of AD are thought to be linked to neuronal dysfunction through the gradual loss of synapses, accumulation of dystrophic neurites that can impair axonal transmission, tau hyperphosphorylation and NFT formation, and eventually the death of neurons. It is unclear whether any of these processes can be halted or reversed, so an appealing therapeutic approach for AD, especially prevention, is neuroprotection- boosting the ability of neurons to resist the toxic effects of amyloid and tau and maintain normal function and viability.[20] Some neurotrophic approaches have included modulation of P75 neurotrophin receptor[21], and AAV mediated delivery of brain-derived neurotrophic factor (BDNF) [22] or nerve growth factor (NGF) [23].

Another protein with potential as a neuroprotective agent in AD and other neurodegenerative diseases is the transcription factor Nuclear factor erythroid 2-related factor 2 (Nrf2, encoded by *Nfe2l2* gene) [24], which regulates the expression of many antioxidant, anti-inflammatory genes [25–29]. Nrf2 is regulated primarily at the protein level through phosphorylation and proteasomal degradation. Kelch-like ECH-associated protein 1 (Keap1), a redox sensor with many cysteines, binds to Nrf2, and interacts with Cullin3 to promote Nrf2 ubiquitination and degradation in the proteosome. When reactive oxygen species accumulate, oxidation of cysteines in Keap1 leads to the release of Nrf2 allowing it to enter the nucleus, bind to transcription partners, such as Maf proteins, and directly regulate target genes via the anti-oxidant response element (ARE) [26, 29]. Regulation of Nrf2 can also occur via phosphorylation, by kinases such as Protein Kinase C (PKC), Glycogen Synthase Kinase 3 (GSK3) PERK, MAPK and CDK5 [30], many of which are dysregulated in AD.

AD brains are marked by increased lipid peroxidation and protein oxidation, leading to the hypothesis that oxidative damage from reactive oxygen species (ROS) contributes to AD onset and progression [31, 32]. There is conflicting evidence regarding Nrf2 in AD patient brains, with some groups reporting reduced mRNA and protein [33, 34], while others report an increase [35]. This may be due to the regions and cells types assessed, with hippocampal neurons having reduced mRNA and protein and reduced nuclear localization [34] and astrocytes having elevated Nrf2 [35].

In mouse models of AD, various studies administering compounds such as sulforaphane that elevate Nrf2 levels have shown cognitive improvements and decreased amyloid and tau pathology [33, 36, 37], while amyloid model mice lacking Nrf2 expression show worsened phenotypes for cognitive function and amyloid deposition [33, 38]. Nrf2 downregulates both the amyloid and tau pathways through different mechanisms. Nrf2 binding to the ARE sequences in Beta-site Amyloid precursor protein (APP) Cleaving Enzyme 1 (BACE1) gene promoter decreases BACE1 expression, while Nrf2 gene disruption elevates BACE1 mRNA and protein in mice [33]. Since BACE1 cleavage of APP is the rate limiting step in Aβ generation, elevated Nrf2 activity could lower amyloid levels. Conversely, Nrf2^-/-^ mice have increased levels of pathological forms of tau associated with AD, such as hyperphosphorylated and sarkosyl insoluble tau and total tau due to reduced autophagic degradation of tau, while compounds that elevated Nrf2 activity increased tau turnover and decreased tau pathological accumulations [39].

In all these studies, the upregulation of Nrf2 through Nrf2-stimulating compounds or reduction through constitutive Nrf2 gene knockout affected all cell types in the brain, as well as peripheral tissue, so the cell types, e.g., microglia, astrocytes, neurons, oligodendrocytes, or some combination, responsible for Nrf2-associated phenotypic changes were unclear. Astrocytic overexpression of Nrf2 is protective in mouse models of AD, Parkison’s, ALS, cerebral hypoperfusion and other neurodegenerative diseases.[40–44] In AD amyloid model APP/PS1 mice and tau model MAPT^P301S^ mice, Nrf2 target genes are upregulated in astrocytes where they act as an adaptive stress response; overexpressing Nrf2 in astrocytes of APP/PS1 and MAPT^P301S^ transgenic mice improved the phenotypes in both models, reducing amyloid burden, tau pathology and synaptic loss [35]. Additionally, RNA sequencing indicates that astrocytic Nrf2 overexpression is neuroprotective by rescuing global transcriptional perturbations in APP/PS1 and MAPT^P301S^ models [35].

While Nrf2 is most highly expressed in astrocytes and microglia in the brain, it is also made in neurons [45], in which data support a protective role for Nrf2 as well. In mouse models of retinitis pigmentosa and macular degeneration, neuronal AAV-mediated Nrf2 overexpression preserved photoreceptors and retinal thickness and led to greater visual function, accompanied by a markedly reduced level of free radicals in the tissue, consistent with upregulation of anti-oxidation enzymes [46–48].

Lentiviral overexpression of Nrf2 in hippocampal neurons of APP/PS1 mice resulted in reduced astrogliosis, improved performance in Morris Water Maze, but no reduction in plaque pathology [49]. In vitro, overexpression of Nrf2 protects neural progenitor cells against Ab toxicity [50]. Additionally, KEAP^-/-^ primary neurons, which have high levels of Nrf2 nuclear translocation, are more resistant to oxidative stresses than KEAP^+/+^ neurons that have less Nrf2 activity, supporting a role for neuronal Nrf2 in cell autonomous protection [51].

Based on these data, overexpression or induction of Nrf2 in neurons could be an effective way to slow or prevent AD progression or onset. In addition, it has been reported that Nrf2 expression in neural progenitor cells is important for neurogenesis,[50] further suggesting that increased Nrf2 in neurons could protect against neurodegeneration by preserving neurogenesis, which is compromised in AD [52, 53].

To gain insight into the molecular mechanism of Nrf2 in AD neuroprotection specifically in neurons, here we used AAV to overexpress Nrf2 in neurons of 5XFAD amyloid pathology model mice. While neuronal overexpression of Nrf2 did not prevent neuronal loss or decrease neuroinflammation or amyloid deposition, we found that the formation of swollen axonal dystrophic neurites around plaques was reduced, along with proteins that accumulate in them, such as BACE1, LAMP1 and phospho-tau 181. Analysis of proteins involved in autophagy suggested this process was unlikely to be involved in reduced dystrophic neurite formation, while RNA sequencing indicated that changes in lipid metabolism and microtubule stability may have been contributing factors. We conclude that Nrf2, while not playing a neuronal cell-autonomous role in reducing amyloid plaques, was effective at reducing dystrophic neurites, which impair action potential conductance [54] and contribute to tau seeding and spreading. Further study of the mechanisms by which Nrf2 protects against dystrophic neurites may lead to novel AD therapeutic strategies. These findings also might suggest a complementary and possibly additive mechanism to that seen in astrocytes, suggesting that expression in both cell types would be more beneficial than expression in either alone.

## Methods

### Mice

5XFAD B6/SJL F1 hybrid on mice were bred in-house by crossing 5XFAD transgenic males to B6/SJL F1 hybrid females, and genotyping of PS1 transgene performed in house as described [55].

Mice were sacrificed by a lethal dose of ketamine/xylazine, followed by transcardial perfusion with 10ml ice cold 1xPBS containing protease and phosphatase inhibitors. One hemibrain was dissected into hippocampus and cortex which were snap-frozen separately. The other hemibrain was drop fixed in 10% buffered formalin overnight at 4°C, transferred to 20% w/v sucrose in 1xPBS for 24 hours, then stored in 30% w/v sucrose in 1xPBS with sodium azide. All animal work was done with the approval of the Northwestern University IACUC, assurance number A3283-01. The number and sex of animals used per experiment is described in detail in the specific methods section of the experiment.

### AAV vector design and preparation and P0 injection of pups

The plasmid of pAAV-Syn-EGFP was acquired from Addgene (catalog #50465), which was deposited by the Bryan Roth Lab at UNC. The cDNA of human Nrf2 (NCBI: NM_006164) was purchased from GeneCopoeia (Rockville, MD; catalog #: T3128). Stbl3 chemically competent E. coli (Thermo Fisher Scientific) was used to transform and amplify all the plasmids. pAAV-Syn-Nrf2 was made by replacing the EGFP sequence of pAAV-Syn-EGFP with the PCR-amplified human Nrf2 cDNA fragments using Gibson Assembly,[56] https://doi.org/10.1101/2020.06.14.150979). The BamHI/EcoRI digestion sites of pAAV-Syn- backbone was destroyed during the cloning process of pAAV-Syn-Nrf2, as the two sites were not included in the overlapping sequences of the 5′- and 3′- end primers for Nrf2 PCR. The full sequence of pAAV-Syn-Nrf2, including the ITR regions, was verified with complete plasmid sequencing technology (MGH CCIB DNA Core). The pAAV-Syn-GFP and -Nrf2 vectors were packaged in AAV serotype 8 capsid using 293 T cells by harvesting the transfected cell culture medium, and they were further purified with iodixanol gradient as previously [57, 58]. The titer of AAV was measured using protein gel methodology by matching the virion protein bands with multiple standard dilutions. The final concentrations of the AAVs were>1 × 10^13^ gc/mL.

5XFAD transgene positive males were crossed to SJL/B6 hybrid females in timed matings to generate transgene negative and positive littermates. On P0, each pup in a litter was cryo-anesthetized and bilaterally injected into the lateral ventricles with 2 µl/hemisphere containing the following amounts of AAV: for “GFP” only, 1×10^9^ viral genomes (VG) syn-GFP; for “Nrf2+GFP” 1×10^9^ VG syn-GFP and 3.33×10^9^ VG syn-Nrf2, and for “Nrf2 high”, 1×10^9^ VG syn-GFP and 1×10^10^ VG syn-Nrf2. All injections were done using a using a 10µl Hamilton syringe with a 30 gauge replaceable needle, as described.[59–61] For each AAV condition, four litters were injected, and four litters were used to generate uninjected controls, for a total of 16 litters. Pups were returned to the mother and aged to 9.5 months.

### Immunofluorescence and image quantification

10% formalin-fixed hemibrains were sectioned at 30μm on a freezing sliding microtome and collected in cryopreserve (30% w/v sucrose, 30% ethylene glycol in 1X PBS) then stored at -20°C. If required, antigen retrieval was performed for 35 minutes at 80°C in 0.1M sodium citrate, in TBS, then allowed to cool for 15 minutes before rinsing well and proceeding with staining. Sections were blocked in 5% normal donkey serum in TBS with 0.25% Triton-X 100 (TBS-T) then primary antibodies added in 1% BSA in TBS-T at 4°C overnight. The following primary antibodies were used: Iba1 (Novus, NB100-1028, 1:300), GFAP (Abcam, ab4674, 1:2000), BACE1 (Abcam, AB108394, 1:300), NeuN (Millipore, Abn91, 1:1000), LAMP1 (Developmental Studies Hybridoma Bank, 1D4B clone, purified antibody, 1:1000), p-tau181 (Cell Signaling, #12885, 1:300), Nrf2 (Santa Cruz, sc518033 (H-6) 1:100 with antigen retrieval), GFP (Abcam Ab5450, 1:1800) Aβ42 (Invitrogen #700254, 1:1000). Following washes, sections were incubated for 2 hours at room temperature with 1:750 dilutions of Alexa Fluor 405, 488, 568 or 647 conjugated donkey anti-mouse, goat, chicken, rat or rabbit secondary antibodies, as appropriate, along with the following stains: 300nM DAPI, and 1:15,000 dilution of 1 mg/ml Thiazine Red (ThR) or MethoxyX04 (MeX) (HelloBio). All secondary antibodies were obtained from ThermoFisher Scientific, except donkey anti-chicken 405, 488 and 647 were from Jackson Immunologicals (West Grove, PA, USA) and donkey anti-rat 568 from Abcam (Waltham, MA USA). Sections were mounted with Prolong Gold (Molecular Probes).

For quantification of BACE1, LAMP1, p-tau181, Iba1 and GFAP three sections between Bregma -1.58mm to Bregma -2.54mm were stained and imaged per mouse. Images were acquired on a Nikon Ti2 Eclipse widefield microscope with a 10x objective, using the NIS Elements software high content method to capture and tile whole sections. All image acquisition settings were maintained the same between treatment groups and genotypes. For image analysis, ROIs were drawn in cortex and hippocampus. Using the General Analysis tool, thresholding was set to distinguish BACE1, LAMP1, Tau p181, Iba1, GFAP, Aβ42, MethoxyXO4, and Thiazine Red positive regions, and calculate total area of ROI, total area positive for each antibody or stain, number of objects positive for each stain, and average size of these objects. Then the percent area covered by each stain from the hippocampal or cortical ROI was calculated.

### Immunoblotting

Snap frozen cortices were homogenized in 1000μl of T-PER Tissue Protein Extraction Reagent (ThermoFisher), supplemented with protease inhibitors (Calbiochem) and Halt Phosphatase Inhibitor Cocktail (Thermo Scientific). Protein concentration was quantified using BCA Assay (Pierce). 20μg of protein was boiled 10min in 1X LDS sample loading buffer, then separated on 4–12% NuPage Bis-Tris Bolt gels (FisherScientific) in MES buffer (FisherScientific). Protein was transferred to nitrocellulose membrane using BioRad Trans Blot Turbo Transfer System, then stained with 0.1% Ponceau in 5% w/v acetic acid and imaged. Membrane was rinsed well, blocked, then probed with the following primary antibodies: anti-APP (6E10, BioLegend 803001, 1:2000), anti-GFP (Abcam ab5450 1:2000), anti-GAPDH (Cell Signaling, #2118, 1:10,000), anti-BACE1 (clone 3D5, Vassar Lab, 1:1000), anti-Nrf2 (clone H-6, Santa Cruz, sc518033, 1:1000), p62 (Cell Signaling, #23214, 1:1,000), LC3BI/II (Cell Signaling, #3868, 1:2,000) ubiquitin (Sigma Aldrich, # 05-944, clone P4D1-A11), anti-Kif3C (Novus Bio-Techne # NBP2-08129, 1:1000) followed by HRP-conjugated anti-mouse, anti-rabbit, or anti-goat secondary antibody (Vector Laboratories 1:10,000). 5% milk was used as a blocking agent, except for Nrf2 antibody, SuperBlock Blocking Buffer (Thermo Fisher Scientific) was used. Blots were visualized using chemiluminescence (BioRad Clarity or ThermoFisher Scientific Pico or West Femto), band intensities measured using a BioRad ChemiDoc Touch Imaging System, then quantified with ImageLab software (BioRad). Signal intensities were normalized to that of GAPDH. Statistical analyses were performed as described below.

### mRNA isolation and bulk mRNA sequencing

For bulk RNA sequencing, RNA was extracted from the hippocampus of the following AAV injected mice described above: non-Tg GFP only (n=6, 3F, 3M), non-Tg Nrf2+GFP (n=6, 3F, 3M), 5XFAD GFP only (n=6, 3F, 3M) and 5XFAD Nrf2+GFP (n=6, 3F, 3M) using the RNAeasy midi kit (Qiagen). The stranded mRNA-seq was conducted in the Northwestern University NUSeq Core Facility. Briefly, total RNA examples were checked for quality using RNA Integrity numbers (RINs) generated from Agilent Bioanalyzer 2100. RNA quantity was determined with Qubit fluorometer. The Illumina Stranded mRNA Library Preparation Kit was used to prepare sequencing libraries from 500 ng of high-quality RNA samples (RIN>7). The Kit procedure was performed without modifications. This procedure includes mRNA purification and fragmentation, cDNA synthesis, 3’ end adenylation, Illumina adapter ligation, library PCR amplification and validation. The Illumina NovaSeq 6000 sequencer was used to sequence the libraries with the production of single-end, 50 bp reads at the depth of 20-25 M reads per sample. The quality of reads, in FASTQ format, was evaluated using FastQC. Reads were trimmed to remove Illumina adapters from the 3′ ends using cutadapt [62]. Trimmed reads were aligned to the Mus musculus genome (mm39) using STAR (https://github.com/alexdobin/STAR). Read counts for each gene were calculated using htseq-count [63] in conjunction with a gene annotation file for mm39 obtained from Ensembl (http://useast.ensembl.org/index.html). Normalization and differential expression were calculated using DESeq2, which uses the Wald test (https://bioconductor.org/packages/release/bioc/html/DESeq2.html). The cutoff for determining significant DEGs was an FDR-adjusted P value of less than 0.05 using the BH method. Pathway analysis was performed using Metascape.

### Statistics

Student’s two-tailed t-test, one way and two-way ANOVA with Sidak’s multiple comparisons test, and linear regression with Pearson’s correlation were done using InStat software (GraphPad Software, Inc., San Diego, CA) to compare means of genotypes, and treatment groups. * 0.05 > p > 0.01 ** 0.01 > p > 0.001 *** 0.001 > p > 0.0001; Error bars = S.E.M.

## Results

### Neuronal overexpression of Nrf2 reduces BACE1

To assess the cell-autonomous effects of neuronal Nrf2 for neuroprotection, we overexpressed Nrf2 directly in neurons of the 5XFAD mouse model of amyloid pathology. Based on previous work, we hypothesized that neuronal Nrf2 overexpression would decrease BACE1 levels and Aβ generation and neuronal loss. Adeno-associated virus 8 (AAV8) expressing human Nrf2 from the neuron-specific synapsin promoter was bilaterally injected into the lateral ventricles of 5XFAD and non-transgenic control mouse pups at post-natal day 0, resulting in lifetime expression of the episomal Nrf2 gene in neurons (Fig 1A) [59]. Mice received one of 4 types of AAV injections: 1) syn-GFP alone, 2×10^9^ VG (GFP only), 2) syn-GFP, 2×10^9^ VG, plus syn-Nrf2, 2.2×10^10^ VG (Nrf2 low), 3) syn-GFP, 2×10^9^ VG, plus syn-Nrf2, 6.6×10^10^ VG (Nrf2 high), or 4) no injection (uninj). At 9.5 months, mice were perfused with PBS and brains collected to process protein, mRNA, or immunofluorescence microscopy analyses. Early analysis by immunofluorescence microscopy indicated that Nrf2 high injection had detrimental effects, including increased plaque load, increased astrocyte and microglial reactivity, and decreased the area covered by the neuronal marker NeuN, suggesting neuronal death compared to uninjected mice (Supplemental Figure 1) so Nrf2 high injected mice were excluded from further analysis.

**Figure 1:**
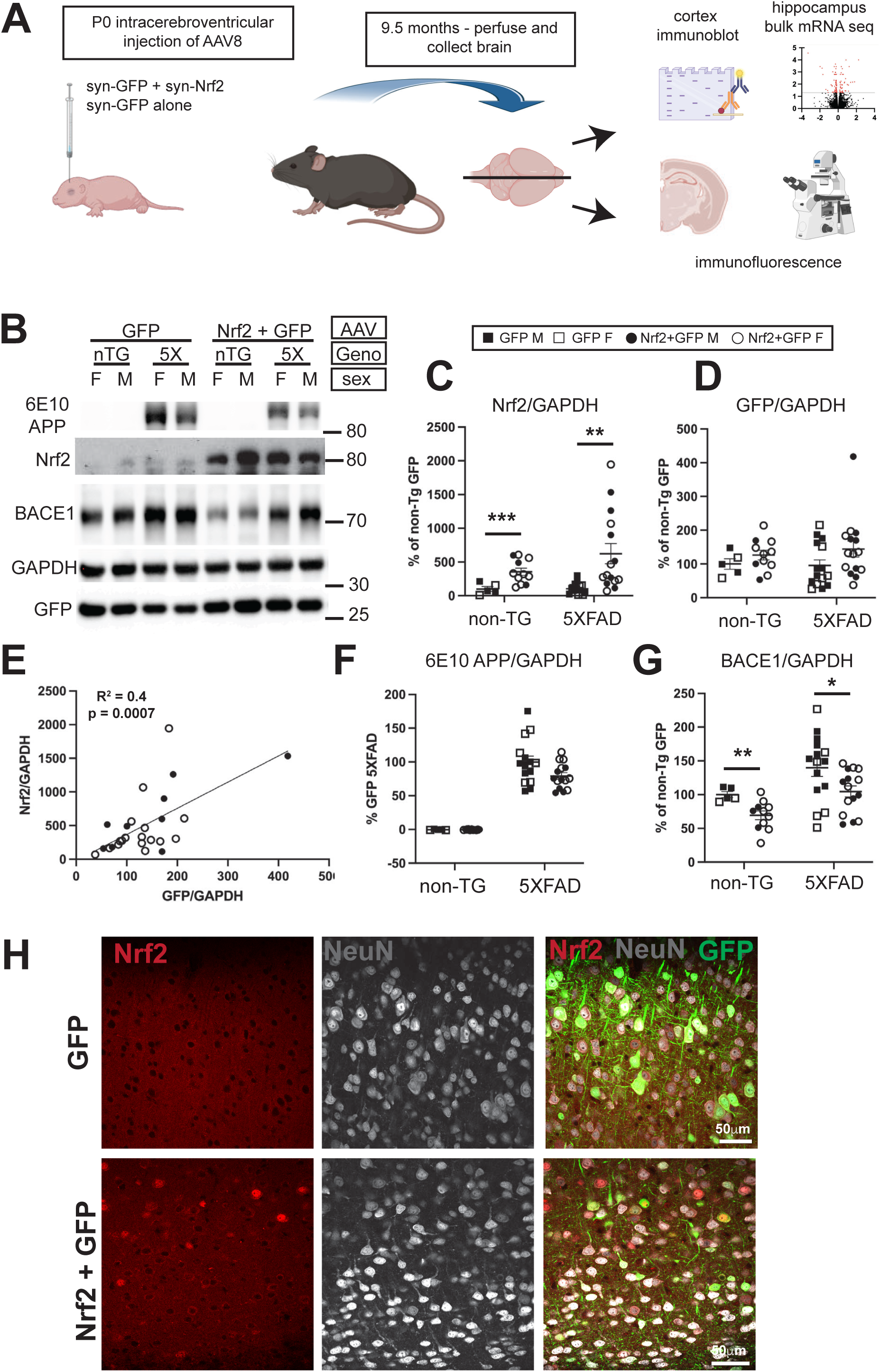
Characterization of Nrf2 overexpression in neurons of 5XFAD mice. (**A**) Schematic diagram of experiment showing AAV injection at P0 followed by aging, harvest at 9.5 months, and analysis by immunoblotting, mRNA sequencing and immunofluorescence microscopy. (**B**) Representative immunoblot images for APP transgenic, Nrf2, GAPDH, and GFP proteins. (**C**) Quantification of Nrf2 protein from immunoblots normalized to GAPDH showing low endogenous levels in GFP only AAV injected mice, but significantly elevated Nrf2 by injection of Nrf2 low AAV (Student’s t-test, Welch correction). (**D**) Quantification of GFP protein in immunoblots normalized to GAPDH is not significantly different between groups. (**E**) Pearsons correlation showing GFP and Nrf2 levels are significantly correlated in mice that received both types of AAV, demonstrating GFP is a surrogate marker of Nrf2 expression. (**F**) Quantification of human APP protein in immunoblots normalized to GAPDH showing transgene-derived APP levels in 5XFAD mice are not significantly affected by Nrf2 overexpression (Student’s t-test). For transgene quantification, the 5XFAD GFP group is set to 100% rather than the non-Tg GFP as in (**D**) and (**G**) since expression of APP/PS1 transgene is not present at all in non-Tg animals. (**G**) Quantification of BACE1 protein in immunoblots normalized to GAPDH showing BACE1 is significantly reduced by Nrf2 overexpression in both 5XFAD and non-transgenic mice (Students t-test) (**H**) Immunofluorescence microscopy images of non-transgenic mouse cortex transduced with Nrf2 low AAV or GFP-only AAV and immunostained for Nrf2, GFP, and NeuN. Note the nuclear localization of Nrf2 in cortical neurons marked by NeuN in the Nrf2+GFP transduced section. Size bar = 50μm. *p<0.05, ** p<0.01, ***p<0.001 Non-TG GFP *n=*5 (2F,3M), non-TG Nrf2+GFP *n=* 11 (8F, 3M), 5XFAD GFP *n=*15 (6F, 9M), 5XFAD GFP+Nrf2 *n=*15 (8F, 7M).

By immunoblot analysis, Nrf2 expression was greatly increased in cortices of Nrf2 low AAV injected mice compared to endogenous Nrf2, which was barely detectable (Fig. 1B, C). GFP expression was similar in Nrf2 low and GFP only groups (Fig. 1B, D) and correlated significantly with Nrf2 levels indicating it is an effective proxy for Nrf2 expression, even though Nrf2 and GFP are not encoded on the same construct but expressed from two distinct co-injected AAVs (Fig 1 E). Nrf2 overexpression did not affect 5XFAD transgene expression, (Fig. 1F). BACE1 levels were decreased in both non-transgenic and 5XFAD mice overexpressing Nrf2 low (Fig. 1G). Confocal microscopy showed Nrf2 expression in neurons of the cortex, identified with neuronal marker NeuN, localizing primarily to the nucleus where it activates gene transcription (Fig. 1H). Together, these results demonstrate that neuronal Nrf2 AAV transduction leads to significant overexpression of Nrf2 that correlates with GFP and causes reduction of BACE1.

### Nrf2 overexpression in 5XFAD mice does not affect plaque load, neuroinflammation, or neuron loss

Next, we analyzed amyloid plaque load, neuroinflammation, and neuronal loss in the cortex and hippocampus of neuronal Nrf2 overexpressing 5XFAD and non-transgenic controls at 9.5 months by widefield immunofluorescence microscopy (representative images in Fig. 2A). Two-way ANOVA was used to determine effect of genotype and AAV. Unexpectedly, there was no significant difference in the percent area covered by amyloid plaques in cortex or hippocampus, as measured by amyloid binding dye MethoxyXO4 (MeXO4) in 5XFAD mice transduced with Nrf2+GFP AAV compared to 5XFAD receiving GFP only AAV, despite reduction in BACE1 (Fig. 2B). Astrocytosis, measured by GFAP immunostaining, and microgliosis, measured by Iba1 immunostaining, were also unaffected by neuronal Nrf2 expression in 5XFAD cortex, while in hippocampus, GFAP and Iba1 immunostainings were reduced in Nrf2+GFP AAV transduced 5XFAD mice compared to GFP-only AAV 5XFAD mice, with the difference reaching statistical significance for GFAP but a non-significant trend for Iba1 (Fig. 2C, D). As expected, genotype significantly affected the area of MethoxyXO4, GFAP and Iba1 in both cortex and hippocampus with all measures being significantly elevated by the 5XFAD genotype, regardless of the type of AAV injected, (Fig. 2B,C,D). In layer 5 of the cortex, neuronal Nrf2 overexpression did not rescue neuronal loss as measured by NeuN-positive immunostaining area in 5XFAD compared to non-transgenic cortex and compared to GFP-only AAV transduction. Although neuronal Nrf2 overexpression did not cause neurodegeneration in 5XFAD mice, we note that it may have promoted neuronal loss as measured by NeuN-positive immunostaining area in the cortex, but not the hippocampus, of non-transgenic animals (Fig. 2E; 2-way ANOVA genotype (5XFAD vs nTG) effect p<0.0001, AAV (Nrf2 vs GFP) effect p=0.004), the significance of which is currently unknown. In hippocampus a small but statistically significant decrease of NeuN positive area in 5XFAD Nrf2+GFP compared to non-Tg Nrf2+GFP was the only significant difference (Fig 2E) (2-way ANOVA, genotype (5XFAD vs nTG) effect p<0.06, AAV (Nrf2 vs GFP) no effect. Overall, these data demonstrate that neuronal Nrf2 overexpression does not reduce amyloid or promote neuronal survival in 5XFAD mice.

**Figure 2:**
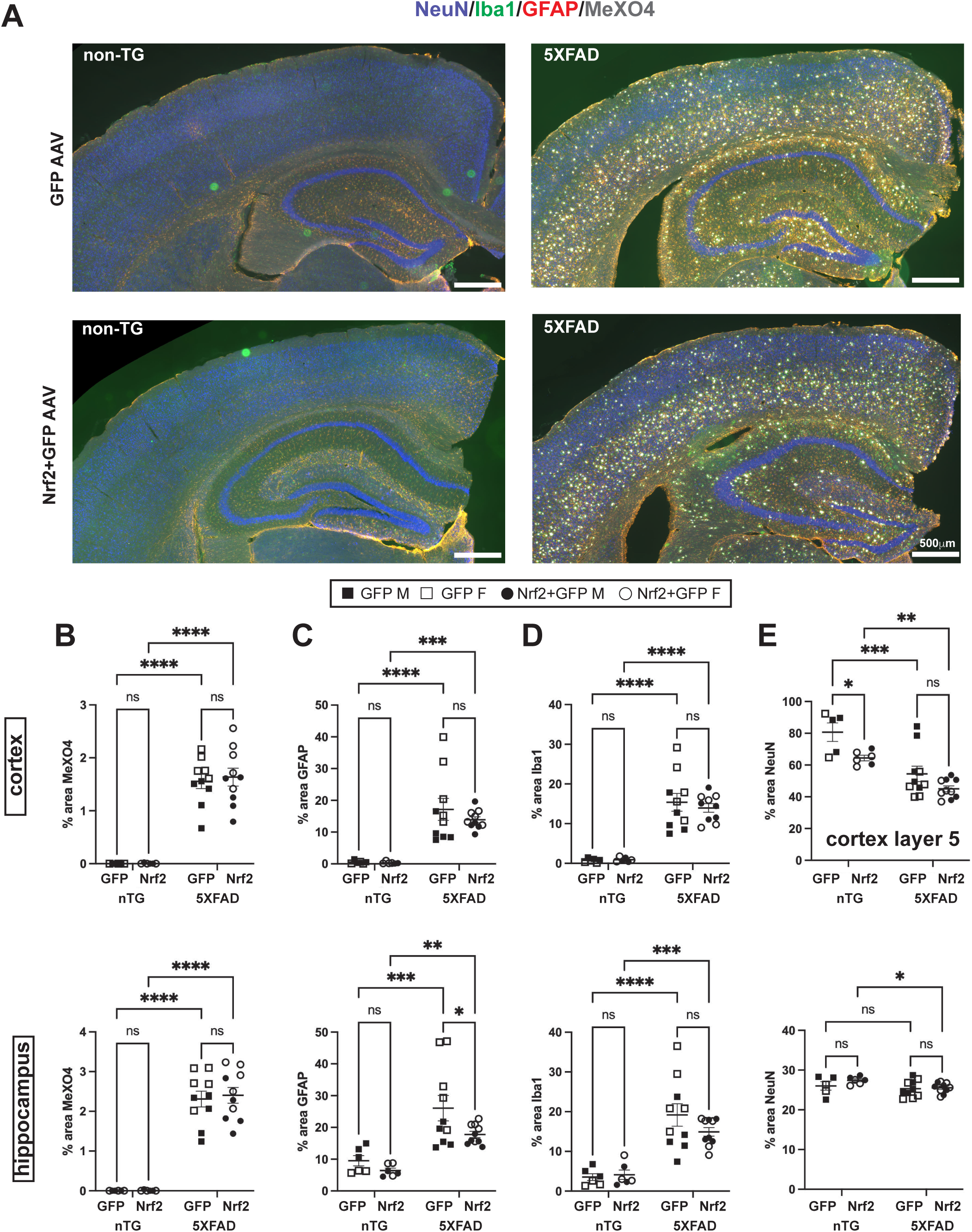
Neuronal Nrf2 overexpression does not affect plaque load, neuroinflammation or neuron loss in 5XFAD mice. (**A**) Representative widefield microscopy of 5XFAD and non-transgenic mouse brain sections immunostained for NeuN (blue, neurons), Iba1 (green, microglia), GFAP (red, astrocytes), and MethoxyXO4 (white, dense core plaques). NB: Images are false colored for viewing. Imaging channels were as follows: MeXO4- 405nm, NeuN - 488nm, GFAP – 568nm, Iba1 – 647nm. Quantification of the percent area in cortex and hippocampus covered by (**B**) amyloid plaques stained with MeXO4 (**C**) astrocytes stained with GFAP (**D**) microglia stained with Iba1 (**E**) neurons stained with NeuN in Layer 5 cortex and hippocampus. 2-way ANOVA with Sidak’s multiple comparisons test. * p<0.05, ** p<0.01, ***p<0.001, **** p<0.0001. Non-TG GFP *n=*5 (3F, 3M), non-TG Nrf2+GFP *n=* 6 (4F, 2M), 5XFAD GFP *n=*10 (5F, 5M), 5XFAD GFP+Nrf2 *n=*10 (5F, 5M). Size bars = 500μm

### Neuronal Nrf2 expression reduces dystrophic neurites around plaques

Another neuropathological feature of AD are peri-plaque dystrophic neurites, which are swollen axons that accumulate proteins such as BACE1, LAMP1, synaptophysin, and other presynaptic proteins, likely due to impaired trafficking and protein degradation [64–67]. Dystrophic neurites (also referred to as axonal spheroids) are detrimental to brain function in multiple ways and are sites of BACE1, APP, and PS1 accumulation and further Aβ generation, [65, 68–72] as well as accumulation of tau phosphorylated at threonine 181,[61] which is a validated CSF and blood biomarker for AD [73, 74]. Dystrophic neurites also impair action potential conduction and inhibit neurotransmission [54], and lack properly formed pre-synaptic densities [71].

To confirm the reduction in BACE1 protein observed by immunoblot (Fig. 1G) and assess BACE1 accumulation in dystrophic neurites, we immunostained brain sections of AAV-transduced mice for BACE1 and Thiazine red to mark plaques (Fig. 3A, B, C). BACE1 is localized prominently in mossy fibers of the dentate gyrus and dystrophic neurites around amyloid plaque cores, as previously reported [55]. As expected, we found that the percent area positive for BACE1 in both cortex and hippocampus was elevated in 5XFAD GFP compared to non-transgenic GFP mice due to BACE1 accumulation in dystrophic neurites. BACE1 was significantly reduced in 5XFAD Nrf2+GFP AAV-transduced compared to 5XFAD GFP-only mice in hippocampus with a trend for reduction in cortex (Fig. 3B, C). To specifically quantify the reduction in BACE1-positive dystrophic neurites in cortex, we normalized the area of BACE1 to amyloid (Thiazine red; ThR) and found the BACE1:ThR ratio was significantly reduced by neuronal Nrf2 overexpression in 5XFAD Nrf2 mice (Fig. 3E). Vesicles in the endo-lysosomal pathway accumulate at high levels in dystrophic neurites, so the protein lysosome associated membrane protein 1 (LAMP1) is widely used as a dystrophic neurite marker [75, 76]. We found a decreased LAMP1:Aβ42 ratio (Fig. 3D, F) further suggesting a reduction of dystrophic neurites. Lastly, we stained for phosphorylated tau at position 181 (p-tau181), also finding a reduction in the ratio of p181 to ThR (Fig. 3D, G). Additionally, the mean size of the halo of dystrophic neurites around the plaque for BACE1 (Fig. 3H) and LAMP1 (Fig. 3I) was reduced as was the size of dystrophic neurites containing p-tau181 (Fig. 3J). Overall, these data suggest that neuronal overexpression of Nrf2 resulted in a decrease in amyloid plaque-associated dystrophic neurites.

**Figure 3:**
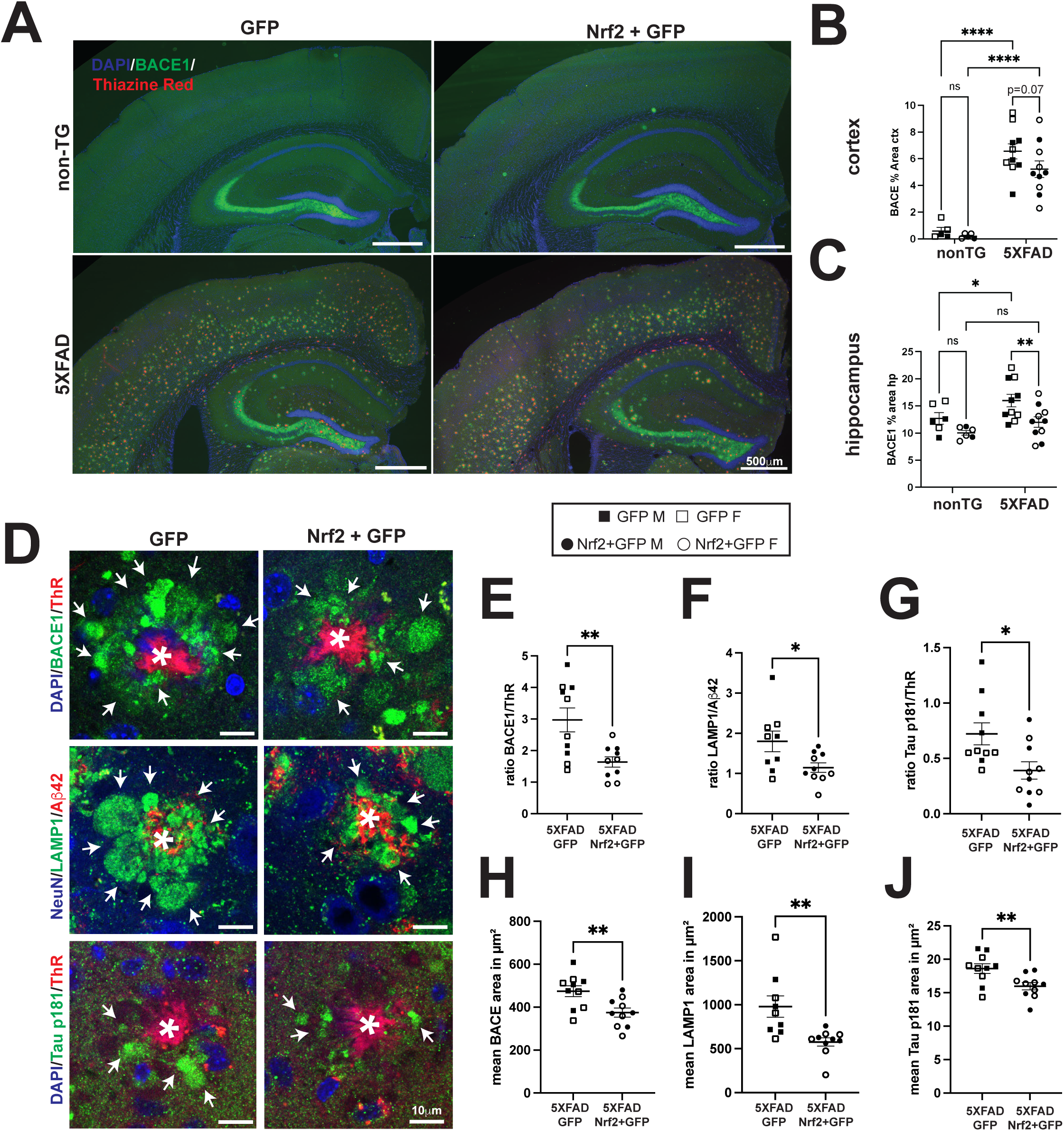
Neuronal Nrf2 overexpression reduces BACE1, LAMP1, p-tau181, and dystrophic neurite halo area in 5XFAD mice. (**A**) Representative brain sections from 5XFAD and non-transgenic (non-TG) mice transduced with Nrf2+GFP or GFP-only AAVs and stained with DAPI (blue), BACE1 (Green) and Thiazine Red (ThR, red) to mark plaques and imaged by widefield immunofluorescence microscopy. Size bar = 500μm (**B**) Percent area covered by BACE immunostaining in 5XFAD cortex is elevated, which has a non-significant downward trend (p=0.07) caused by neuronal Nrf2 overexpression (**C**) Percent area covered by BACE immunostaining in 5XFAD hippocampus is elevated, which is significantly reduced by neuronal Nrf2 overexpression. 2-way ANOVA with Sidak’s multiple comparisons test * p<0.05, ** p<0.01, **** p<0.0001 (**D**) Representative single plane confocal images of amyloid plaques (red, Thiazine red, *= plaque core) surrounded by dystrophic neurites immunostained for BACE1, LAMP1, and p-tau181 (left to right, green) in GFP-only or Nrf2+GFP AAV transduced 5XFAD mice. Note that in the right panel of images from Nrf2 overexpressing mice, dystrophic neurites are smaller, fewer and of less overall plaque coverage than upper panel (arrows). Size bar = 10μm. (**E-G**) Quantification of the ratio of dystrophic neurite proteins (BACE, E; LAMP1, F; tau p181, G) to amyloid plaque core (Thiazine red, E, G; Aβ42, F) in cortex of GFP-only or Nrf2+GFP AAV transduced 5XFAD mice. In all cases, the ratios are reduced in neuronal Nrf2 overexpressing 5XFAD mice, indicating reduced dystrophic neurites for a given amount of amyloid. (**H-J**) Quantification of the mean areas of BACE1, LAMP1 or tau p181 immuno-positive structures around individual amyloid plaques in cortex. For all three proteins, the average areas of immunostaining are reduced in Nrf2 overexpressing mice. (**E-J**) quantification from 10x Ti2 widefield images, using NIS Elements, as described in methods. * p<0.05, ** p<0.01, two tailed t-test. NB: Images are false colored for viewing. Imaging channels were as follows: DAPI, NeuN – 405nm, ThR, Aβ42 – 568nm, BACE1, Aβ42, p181 - 647nm. Non-TG GFP *n=*5 (3F,3M), non-TG Nrf2+GFP *n=* 6 (4F, 2M), 5XFAD GFP *n=*10 (5F, 5M), 5XFAD GFP+Nrf2 *n=*10 (5F, 5M)

### Autophagy proteins are increased in non-Tg mice by neuronal Nrf2 overexpression

Dystrophic neurites are known to accumulate large quantities of autophagic intermediates as well the autophagy protein LC3B [71, 77, 78] and ubiquitinated proteins targeted for degradation by autophagy [79–81]. Studies indicate that early interventions to increase autophagy can improve phenotypes in amyloid pathology mouse models [82, 83]. Nrf2 can promote autophagy by increasing levels of p62, an important autophagy cargo adaptor [84], suggesting that neuronal Nrf2 overexpression may decrease dystrophic neurites by promoting autophagy. To investigate this possibility, we performed immunoblot analysis for key proteins in the autophagy pathway: LC3B-I, which is cleaved to form LC3B-II, an initiator of autophagosome formation; p62, an adaptor protein that recruits ubiquitinated proteins for degradation (Fig. 4A). We also used an anti-ubiquitin antibody to assess levels of ubiquitinated cargo proteins (Fig. 4A). In non-Tg Nrf2 AAV transduced mice, p62 is significantly elevated, indicating an increase in autophagy (Fig 4B), but between 5XFAD Nrf2+GFP AAV transduced and 5XFAD GFP AAV transduced mice p62 is unchanged. The most significant elevation is in the 5XFAD compared to non-transgenic mice, regardless of AAV transduction, congruent with the high levels of autophagy observed in 5XFAD mice. A similar pattern was observed for total LC3B (Fig. 4C) and ubiquitinated proteins (Fig. 4D), with significant increases in 5XFAD mice, and a non-significant trend for elevation in non-transgenic Nrf2+GFP AAV transduced, and no change in 5XFAD Nrf2+GFP AAV transduced. As a measure of autophagic flux we assessed the ratio of LCB-II/LC3B-I, and found again very significant elevations in 5XFAD, while Nrf2 transduction only elevated the ratio in 5XFAD mice (Fig. 4E). Our results suggest that neuronal Nrf2 overexpression may tend to increase autophagy-related proteins, especially in non-transgenic mice, but in nine-month-old 5XFAD mice autophagy induction has reached a ceiling, so Nrf2 mediated enhancement of autophagy is unlikely to play a mechanistic role in Nrf2-mediated dystrophic neurite reduction.

**Figure 4:**
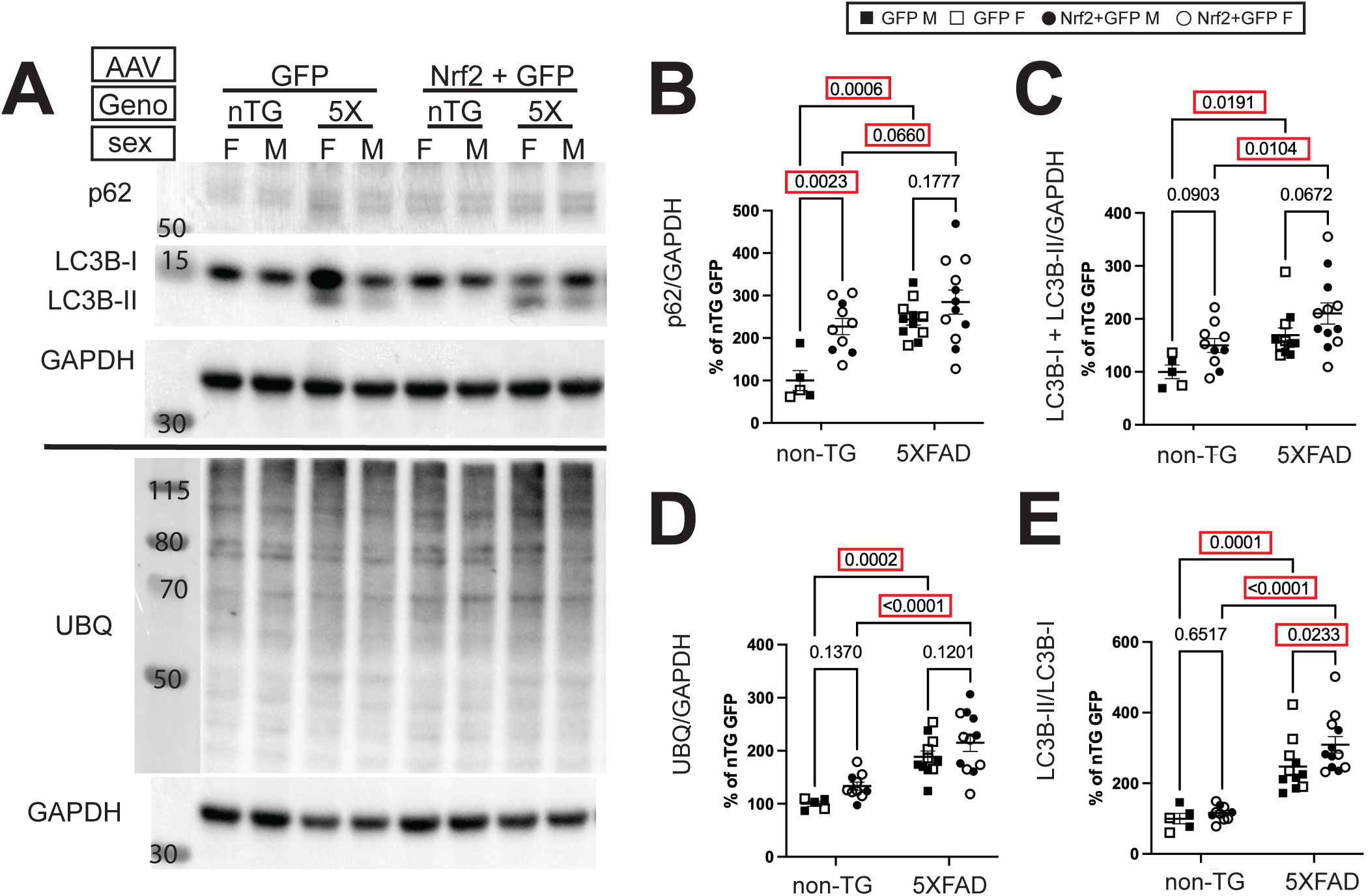
Neuronal Nrf2 overexpression increases autophagy proteins in non-transgenic mice but not 5XFAD mice. (**A**) Representative immunoblots of LC3BI/II, p62, and ubiquitin normalized to GAPDH. (**B**) Autophagy adaptor protein p62 is significantly elevated in 5XFAD genotype, and by Nr2+GFP AAV transduction of non-transgenic mice. (**C**) Total LC3BI and II is also significantly elevated in 5XFAD genotype, with a trend for increase in Nr2+GFP AAV transduced non-transgenic mice. (**D**) ubiquitinated protein shows elevation in 5XFAD genotype, with no significant change resulting from Nrf2 AAV transduction in either 5XFAD or non-transgenic mice (**E**) The ratio between cleaved, active LC3B-II and full length LC3B-I shows elevation in 5XFAD genotype, with a further significant elevation in response to Nrf2+GFP AAV transduction. 2-way ANOVA with Sidak’s multiple comparisons test. These data suggest that autophagy machinery (p62, LC3B-I/II) are upregulated due Nrf2 expression non-transgenic mice, but the already high levels in 5XFAD mice are not further increased. Non-TG GFP *n=*5 (2F,3M), non-TG Nrf2+GFP *n=* 11 (8F, 3M), 5XFAD GFP *n=*15 (6F, 9M), 5XFAD GFP+Nrf2 *n=*15 (8F, 7M).

### Activation of Nrf2 transcriptional networks have differing effects in non-transgenic and 5XFAD mice

To confirm changes to Nrf2 signaling pathways and identify mechanisms that could be responsible for reduced dystrophic neurite formation in Nrf2+GFP AAV transduced 5XFAD mice, we performed bulk mRNA sequencing of 5XFAD hippocampi of Nrf2+GFP AAV transduced mice and GFP AAV transduced controls. As expected, the most significantly upregulated gene in Nrf2+GFP AAV transduced 5XFAD and non-transgenic mice was *Nfe2l2*, which encodes Nrf2 protein (Fig. 5A). Genes downstream of Nrf2 signaling (*Hmox1, Txnrd1, Gsta3, Gsta4, Gstm7*) were also significantly upregulated in Nrf2+GFP AAV transduced mice (green in Fig. 5A, green arrow in Fig. 5B), indicating that AAV-mediated overexpression of Nrf2 had activated its transcriptional pathways. We also compared GFP-only AAV-transduced 5XFAD and non-transgenic mice and found a very large number of upregulated genes (782), many of which were associated with inflammatory or reactive microglia, congruent with other studies of mRNA changes in 5XFAD mice (right panels, Figs. 5A, B) [85–87]. Figure 5 B shows the 50 most significantly changed genes for each comparison, and in the Nrf2+GFP AAV transduced 5XFAD mice, many of these genes are neuronally expressed (*Erbin, Erlin, Postn, Ebf2, Dbi, Bex, Cacul1* - orange arrows) as expected for Nrf2 overexpression driven in neurons. Interestingly, several *Postn* [88], *Erbin* [89], and *Erlin 2* [90], are important in development and neuronal outgrowth. There is not a large overlap of gene expression changes between non-transgenic and 5XFAD Nrf2+GFP AAV transduced mice, comprising only five shared upregulated genes: *Nfel2l* itself, three direct targets, *Txnrd1, Hmox1,* and *Glul*, and *Cdkna1*, which encodes p21, a senescence promoting protein (green arrows), and no shared down regulated genes, indicating that the effects of neuronal Nrf2 overexpression in the context of amyloid are different than in wild type mice.

**Figure 5:**
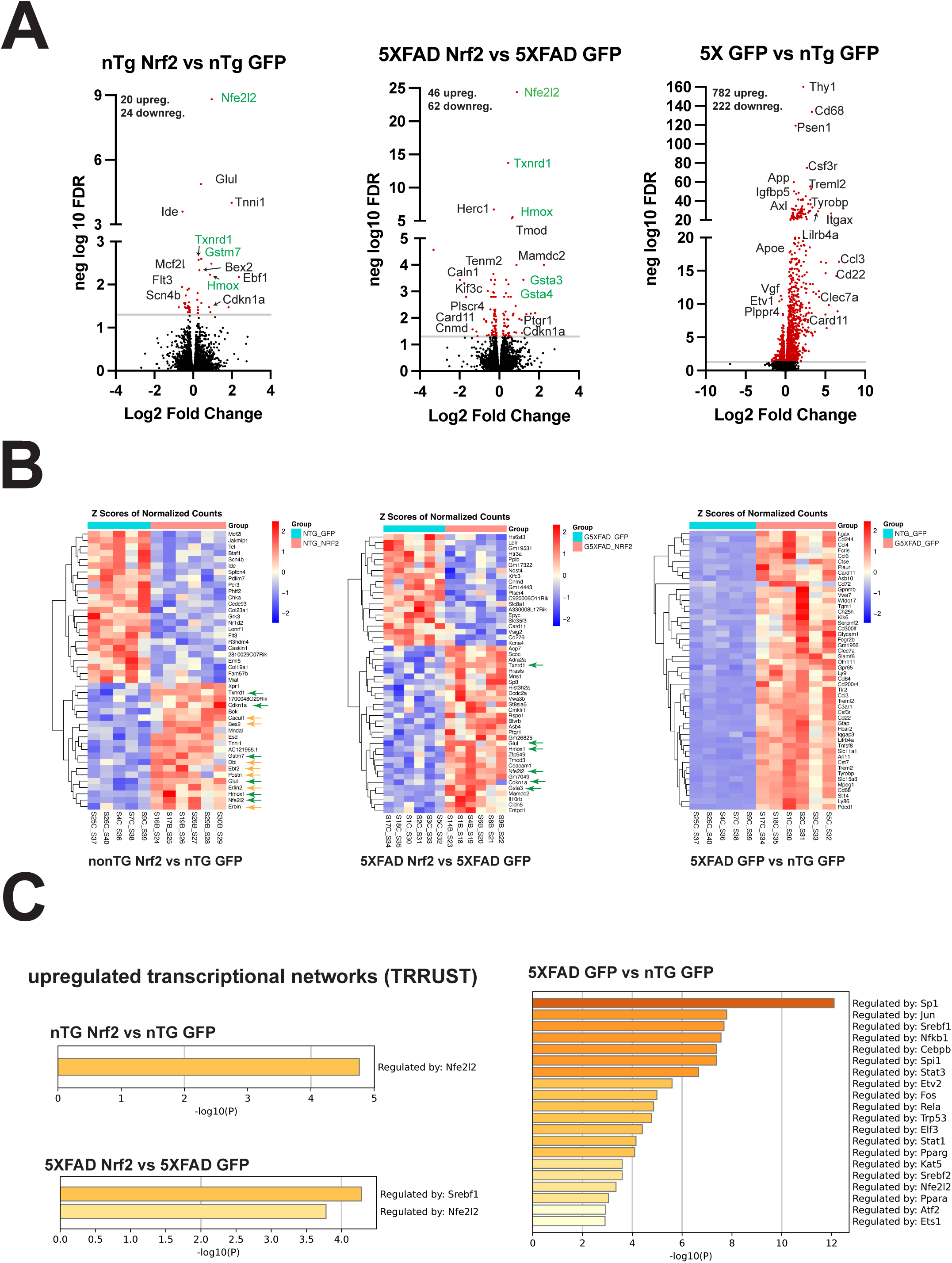
Activation of Nrf2 transcriptional network has differing effects in non-transgenic and 5XFAD mice. (**A**) Volcano plots of differentially expressed genes (DEGs) in hippocampus of (left to right), nTG Nrf2 vs nTG GFP, 5XFAD Nrf2 vs 5XFAD GFP and nTG GFP vs 5XFAD GFP. The first two resulted in smaller number of DEGs than 5XFAD compared to nTG. Green indicates direct Nrf2 (*Nfe2l2*) target genes. 5XFAD is dominated by microglial and astrocytic activation markers. (**B**) Heat maps of 50 most significantly changed DEG, again left to right nTG Nrf2 vs nTG GFP, 5XFAD Nrf2 vs 5XFAD GFP and nTG GFP vs 5XFAD GFP. (**C**) TRRUST (metascape.org) analysis to identify transcription factors associated with the upregulated DEG for each group.

TRRUST analysis of transcriptional network changes (Fig. 5C) shows activation of the *Nfe2l2* network in non-transgenic and 5XFAD Nrf2 mice, and activation of the *Srebf1* transcriptional network in 5XFAD Nrf2 mice. Srebp1 activation may be downstream of Nrf2 since Nrf2 can act with coactivator Brg1 to promote Srebf1 transcription [91]. *Srebf1* encodes the transcription factor Srebp1 which drives expression of genes for fatty acid and cholesterol production [92]. Neuronal expression of Srebp1 functions cell-autonomously to promote dendritic arborization [93], and a polymorphism in Srebp1 has been reported to affect AD risk in APOE4 individuals [94] suggesting a role for Srebp1 function in AD.

In 5XFAD GFP mice compared to non-Tg, many of the activated transcriptional networks drive expression of inflammatory genes in microglia and astrocytes. *Nfe2l2* and *Srebf1* networks are upregulated in 5XFAD mice even in the absence of Nrf2 overexpression, indicating dysregulation of these networks, though in 5XFAD GFP mice the *Nfe2l2* network activation is likely occurring in non-neuronal cells (astrocytes, oligodendrocytes, microglia) where the majority of Nrf2 is expressed [35, 95, 96], so the further increase in 5XFAD Nrf2 compared to 5XFAD GFP is likely to due to the AAV-driven neuronal Nrf2 overexpression.

### Nrf2 overexpression may affect dystrophic neurites through lipid pathways or microtubule stability

We next analyzed the effects of neuronal Nrf2 overexpression in both non-transgenic and 5XFAD mice using Metascape Gene Ontology analysis, which showed upregulation of the hepatocellular carcinoma pathway that is characterized by key Nrf2 genes (*Cdkn1a, Gsta3, Gsta4, Hmox1, Nfe2l2, Txnrd1, Blvrb, Gng12, Entpd*1) involved in antioxidant response, detoxication, NADPH generation, and carbohydrate, lipid and nucleic acid metabolism. In Nrf2+GFP AAV transduced non-transgenic mice, other upregulated processes included response to nutrient levels, apoptotic signaling and regulation of innate immune response, with only one downregulated pathway related to rhythmic process (Fig. 6A). In 5XFAD Nrf2+GFP AAV transduced mice (Fig. 6A), upregulated processes include regulation of angiogenesis *(Cldn5, Glul, Hmox1, Nfe2l2, Ceacam1, Clic4, Rtf2, Cdkn1a*), response to wounding, integration of energy metabolism, and negative regulation of secretion by cells.

**Figure 6:**
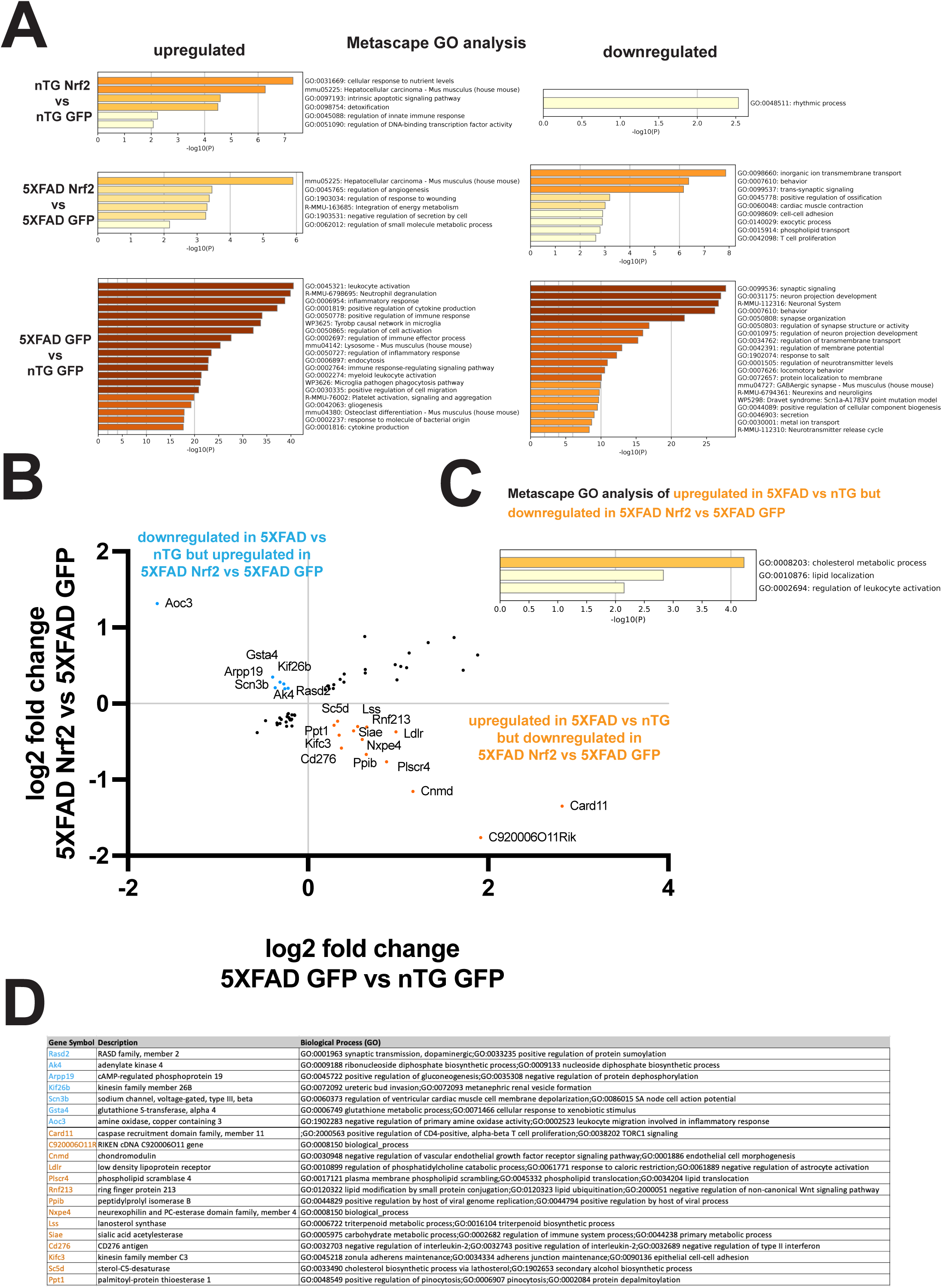
Neuronal Nrf2 overexpression may decrease dystrophic neurites through alterations to lipid pathways or microtubule stability. (**A**) Metascape gene Ontology (GO) of upregulated (left) and downregulated (right) processes in nTG Nrf2 vs nTG GFP, 5XFAD Nrf2 vs 5XFAD GFP and 5XFAD GFP vs nTG GFP. (**B**) Correlation of log2 fold in 5XFAD GFP vs nTG GFP with log2 fold change in 5XFAD Nrf2 vs 5XFAD GFP. Most interesting are those upregulated in 5XFAD GFP vs nTG GFP and downregulated in 5XFAD Nrf2 vs 5XFAD GFP (orange points) or downregulated 5XFAD GFP vs nTG GFP and upregulated in 5XFAD Nrf2 vs 5XFAD GFP (blue points), as they could be showing reversion of 5XFAD changes to the baseline state. (**C**) Metascape Gene Ontology analysis of upregulated in 5XFAD GFP vs nTG GFP and downregulated in 5XFAD Nrf2 vs 5XFAD GFP (orange points) showed changes to cholesterol metabolism (*Ldlr, Lss, Sc5d, Ppt1*) and lipid localization (*Ldlr, Plscr4, Rnf213*). (**D**) List of genes from (**B**) with protein name and GO functional information.

In 5XFAD Nrf2+GFP AAV transduced mice (Fig. 6A), several downregulated processes have neuronal components, such as inorganic ion transmembrane transport (*Gabrb2, Gabrg2, Hcn1, Htr3a, Kcna4, Kcnc3, Scn1a, Slc8a1,Slc24a2, Kcnq3, Scn2a, Kcnh7, Ptprd, Card11*), behavior/memory/cognition (*Camk4, Etv1, Gabrg2, Hcn1, Ldlr, Scn1a, Fezf2, Sorcs3, Slc24a2, Synj1, Kcnq3, Scn2a, Adgrl3, Kcnc3, Ptprd, Prepl*), trans-synaptic signaling (*Camk4, Gabrb2, Gabrg2, Htr3a, Ptprd, Tenm2, Synj1, Kcnq3, Dmxl2, Cadps2, Adgrl3, Hcn1, Slc8a1*), exocytic process (*Exoc1, Synj1, Cadps2, Rasgrp1, Slc8a1, Cyb5r4, Celsr2, Kcnc3, Prepl*) and phospholipid transport (*Ldlr, Atp11b, Plscr4*).

In 5XFAD (Fig. 6A), upregulated processes are highly inflammatory including leukocyte activation, inflammatory response, positive regulation of cytokine production, Tyrobp causal network in microglia, lysosome, endocytosis, and cytokine production, in agreement of other RNA analyses of aged 5XFAD mice [85–87]. Also congruent with other work, downregulated processes included synaptic signaling, neuron projection development, behavior, synapse organization, regulation of neurotransmitter levels, and gabaergic synapse, indicating loss of neuronal function due to synapse and neuronal loss [85–87].

To determine which gene expression changes might be playing a role in the reduction of dystrophic neurites, we looked for genes upregulated in 5XFAD GFP compared to non-transgenic GFP that were downregulated in 5XFAD Nrf2 compared to 5XFAD GFP (14 genes, orange in Fig. 6B) or genes downregulated in 5XFAD GFP compared to non-transgenic GFP that were upregulated in 5XFAD Nrf2 compared to 5XFAD GFP (7 genes, in blue in Fig. 6B), indicating a return to a more homeostatic level in the presence of Nrf2 overexpression. Metascape GO analysis of these 14 genes increased in 5XFAD that were decreased in 5XFAD Nrf2 found upregulation of cholesterol metabolic process (*Ldlr, Lss, Sc5d, Ppt1*) and lipid localization (*Ldlr, Plscr4, Rnf213*), as well as leukocyte activation (*Ldlr, Cd276, Card11*), suggesting changes to lipid handling and immune pathways in 5XFAD Nrf2+GFP AAV transduced compared to 5XFAD GFP AAV transduced (Fig. 6C). Recent work shows accumulation of neutral lipid droplets in proinflammatory plaque-associated microglia [97, 98] and in dystrophic neurites themselves [99], suggesting that altered lipid handling or membrane composition could affect dystrophic neurite formation. Interestingly, the *Srebf1* transcriptional network, which promotes fatty acid and cholesterol production, is upregulated in 5XFAD GFP AAV transduced mice compared to non-transgenic GFP AAV transduced mice, and then further upregulated in the 5XFAD Nrf2+GFP AAV transduced mice compared to 5XFAD GFP AAV transduced mice. Increased fatty acid and cholesterol production may be an adaptive response to resist membrane damage by amyloid that is further promoted by neuronal Nrf2 overexpression.

Two other genes that changed in opposite directions between 5XFAD GFP and non-transgenic and 5XFAD Nrf2 and 5XFAD GFP, *Kif3c* and *Kif26b*, are of high interest because they encode microtubule interacting proteins. Loss of microtubule stability is associated with development of axonal dystrophies due to impaired vesicle transport [65, 100, 101] and microtubule stabilizers are being investigated as a therapeutic approach to protect against dystrophic neurite formation and tau pathology [102, 103].

Kif3c is a microtubule destabilizing protein [104, 105] and is upregulated in 5XFAD mice but decreased by Nrf2+GFP AAV transduction, suggesting improved microtubule stability after neuronal Nrf2 overexpression. *Kif26b* mRNA shows the opposite expression patterns, downregulated in 5XFAD and upregulated by Nrf2 expression, and has the opposite biological effect of Kif3c, serving to stabilize microtubules [106], again suggesting improved microtubule stability after Nrf2 overexpression. A mis-sense mutation, G546S, in Kif26B is associated with microcephaly and pontocerebellar hypoplasia in humans, indicating a key role in neuronal development and function [107]. Additionally, Kif26b is reported to be a protective rare variant associated protein in the AD vulnerable population of somatostatin inhibitory neurons, suggesting a potentially protective effect in AD [108].

Because of the clear connection between microtubule stability and dystrophic neurite formation, we attempted to confirm the changes in Kif26b and Kif3c at the protein level. While we were unable to detect Kif26b via immunoblot analysis, we confirmed that Kif3c protein levels were increased in 5XFAD GFP compared to non-transgenic GFP mice and strongly decreased in 5XFAD Nrf2 compared to 5XFAD GFP mice (Fig. 7A). These results support the hypothesis that Nrf2-mediated reduction in Kif3c contributes to the reduction of dystrophic neurites in 5XFAD mice via microtubule stabilization.

**Figure 7:**
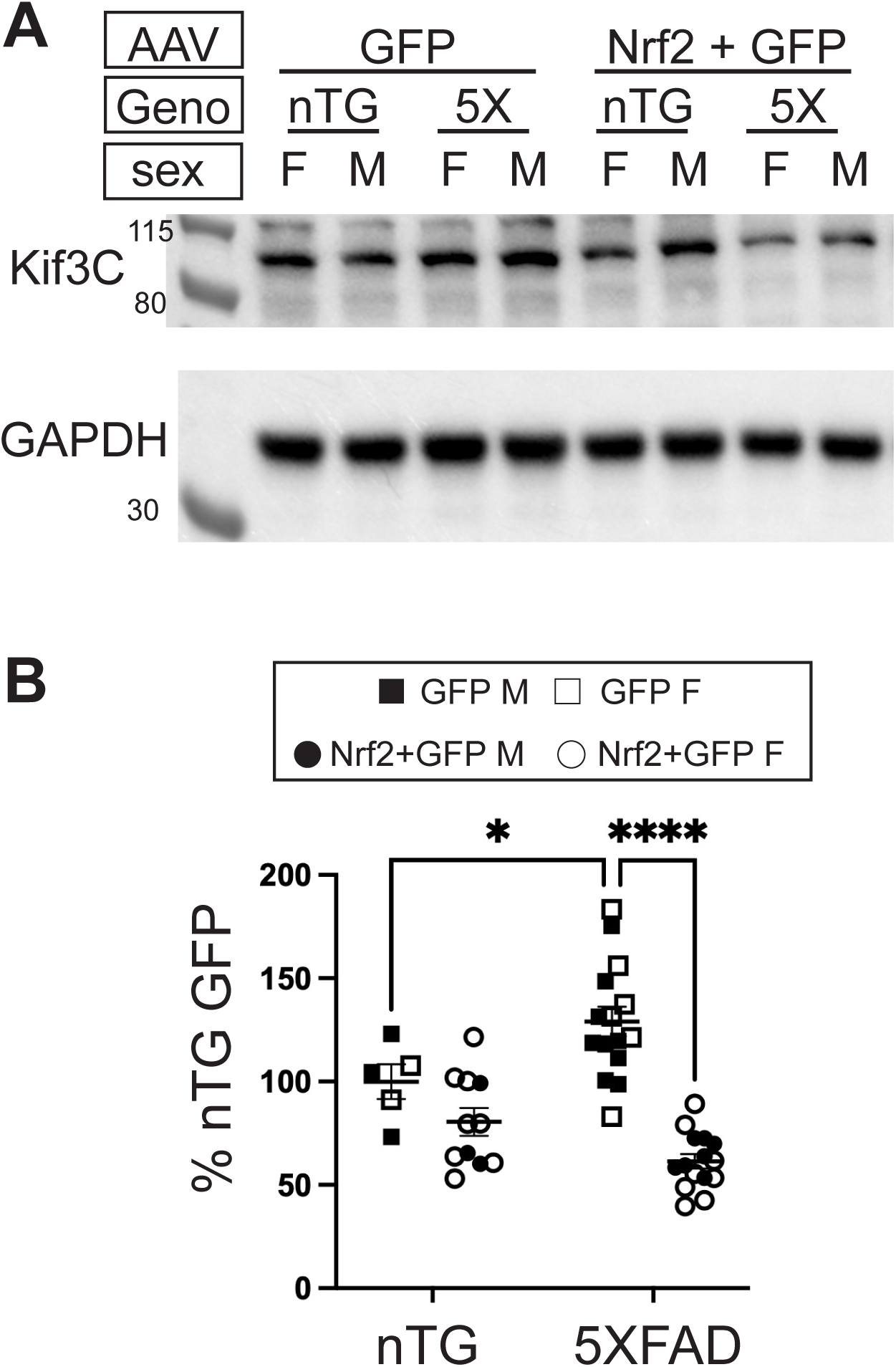
Kif3c protein is elevated in 5XFAD mice and reduced by neuronal Nrf2 overexpression. (**A**) Representative immunoblot analysis of Kif3C and GAPDH proteins from cortical homogenates of 5XFAD and non-transgenic mice transduced with GFP-only or Nrf2+GFP AAVs. (**B**) Kif3C immunoblot signals were normalized to GAPDH and represented as percent of non-transgenic GFP-only control. Quantification shows elevation of Kif3C protein in 5XFAD GFP mice compared to non-Tg GFP and reduction of Kif3C in 5XFAD Nrf2 compared to 5XFAD GFP mice, indicating that Kif3C protein levels follow the same pattern of change as Kif3C mRNA levels. 2-way ANOVA with Sidak’s multiple comparisons test, * p<0.05, **** p<0.0001 Non-TG GFP *n=*5 (2F,3M), non-TG Nrf2+GFP *n=* 11 (8F, 3M), 5XFAD GFP *n=*15 (6F, 9M), 5XFAD GFP+Nrf2 *n=*15 (8F, 7M).

## Discussion

In this study we have demonstrated that Nrf2 can be successfully overexpressed specifically in neurons of non-transgenic and 5XFAD mice after P0 injections of AAV. While we observed neuronal loss for unknown reasons at high levels of Nrf2 AAV, at lower titers we found that AAV overexpression of Nrf2 in neurons protected against dystrophic neurite formation. We confirmed that Nrf2 decreases BACE1 levels but does not reduce the frequency of dense core plaques in 5XFAD mice at nine months. Correspondingly, the activation of microglia and astrocytes are not changed by neuronal Nrf2 overexpression. Interestingly, we found in 5XFAD mice that neuronal Nrf2 overexpression reduced the amount of neuritic dystrophy around plaques, as measured by accumulation of LAMP1, BACE1 and phospho-tau 181. This results opens the possibility that neuronal Nrf2 could have beneficial effects on AD pathology and progression, since high levels of dystrophic neurites are associated with greater seeding and spreading of tau pathology [16, 17].

We had hypothesized that in addition to promoting neuronal survival, Nrf2 overexpression would decrease amyloid load by downregulating BACE1, as described in the literature [33]. On the contrary, we observed that despite the reductions in BACE1, there was no decrease in plaque load. This may be due to the presence of the Swedish mutation in the APP transgene carried by the 5XFAD mice, which enhances the rate of BACE1 cleavage of APP [109]. We have previously observed that in 5XFAD mice, even 50% reduction of BACE1 levels do not reduce Aβ generation [110]. This may be a similar situation, where even in the context of ∼30% BACE reduction, it is still in excess of APP and can cleave all available substrate. It is also possible, that at an earlier timepoint in amyloid deposition, this degree of BACE1 reduction may reduce amyloid deposition, but with time, plaque accumulation equalizes across treatment groups.

We investigated effects of Nrf2 overexpression on the autophagy pathway since this process is in part regulated by Nrf2 and plays a role in dystrophic neurite formation. While we did find that Nrf2 overexpression increased p62 levels in non-transgenic mice, the effects on 5XFAD mice were much less noticeable since all aspects of autophagy we assessed, p62, total LC3B, ubiquitinated cargo and the ratio of LC3B-II:LC3BI were already significantly elevated in the 5XFAD compared to non-transgenic mice. Our experiments did not specifically address autophagic flux, but there is data suggesting flux is blocked in amyloid model mice due to lysosomal deficiency [111] so it may be that promoting initiation of autophagy through Nrf2-mediated p62 increase is not sufficient to improve protein clearance.

Another unexpected finding was the relatively small number of genes altered at the transcriptional level in by Nrf2 overexpression in the non-Tg and 5XFAD animals. While there were key genes in the Nrf2 pathway, it suggests that perhaps with chronic high levels of overexpression the cells adapt by downregulating the Nrf2 pathway, perhaps by upregulating KEAP1 or other proteins that modulate Nrf2 activity. It also may be due to the fact that only neurons were overexpressing Nrf2, so these cell-type specific transcriptional changes were dampened by the lack of change in mRNA from astrocytes, microglia, oligodendrocytes and other cell types that were also present in the bulk analysis. While P0 injection of AAV8 results in wide-spread gene expression throughout the brain, there are some neurons that will not be transduced or do not express from human synapsin promoter we used, which could further dilute the bulk mRNA signal. Additionally, it has been reported that in mice with *Nfe2l2* gene deletion in specific cell types of the brain (astrocytes, microglia) only about 60 transcripts are significantly changed [112], similar to the reported 40 (nonTg) or 110 (5XFAD) DEGs in this study. Single nuclear RNA sequencing focusing on neurons could describe in more detail the Nrf2-induced transcriptional changes to help shed light on the mechanism responsible for decreased dystrophic neurites.

One of the most interesting observations from this study is that Nrf2 overexpression results in a decrease of dystrophic neurites even without reductions in amyloid plaques, providing new insights into mechanisms governing dystrophic neurite formation. Dystrophic neurite reduction cannot be attributed to neuronal loss as 5XFAD Nrf2 and 5XFAD GFP have no significant difference in NeuN immunostaining (Fig. 2E). From our bulk RNA sequencing, two mechanisms that could play a role in rescuing dystrophic neurites emerged: lipid processing and microtubule stability. Cholesterol and other lipid alterations have been observed in AD [113, 114] and multiple AD risk genes such as APOE, TREM2, CLU, ABCA1, ABCA7, SORL, PICALM, BIN1 are involved in lipid handling in the brain (reviewed in [115]). Further studies to describe the specific lipid changes found in neuronal Nrf2 overexpressing mice could help determine how these changes protect against dystrophic neurite formation, perhaps though increasing lysosomal efficiency [116], or making the neuronal membrane more resistant to amyloid-mediated damage [117]. Increasing stability of microtubules is associated with decreased dystrophic neurites [100, 101, 103], so we predict that overexpressing the stabilizing protein Kif26b and/or reducing the destabilizer Kif3c would protect against neuritic dystrophy and subsequent tau spread. Future work will address these questions, shedding further light on the transcriptional mechanisms of Nrf2 and understanding of dystrophic neurite formation.

### Limitations

Our study has several caveats that limit our conclusion that direct neuronal expression of Nrf2 does not prevent neuronal death in 5XFAD cortex. We cannot rule out the possibility that an even lower level of Nrf2 expression in neurons might promote neuronal survival, as the lowest Nrf2+GFP AAV transduction we used resulted in much higher protein levels (Fig. 1C, ∼250-600x) than baseline, which could still have negative consequences. Nrf2 is developmentally regulated in the brain, with highest levels occurring during neuronal differentiation, and minimal levels detected in adult neurons, likely due to a repressive chromatin structure [95, 96], so it is possible that maintaining active Nrf2 signaling pathways throughout life may have adversely affected neuronal differentiation or maintenance of the differentiated state. This could explain why the very high levels of Nrf2 in the Nrf2 high AAV transduced mice caused loss of NeuN staining (Supp. Fig 1E). Additionally, since we did not look at other timepoints, it is possible that the lower Nrf2 overexpression could have more of an effect later as neuronal loss accelerates between 9 and 12 months [55].

## Conclusions

While previous work indicated beneficial effects of Nrf2 expression globally or in astrocytes in the context of Alzheimer’s and other neurodegenerative disease, the data we present show a role for Nrf2 in protecting against a specific neuronal pathology, dystrophic neurites, that have adverse effects such as impairing action potential conductance and contribute to tau seeding and spreading. Their reduction by neuronal Nrf2 overexpression may protect neurons against these pathologic changes, suggesting a beneficial effect of Nrf2 in neurons. Additionally, further study of the pathways by which Nrf2 reduces dystrophic neurites may lead to novel therapeutic strategies that can limit neuritic damage, dystrophic neurites, and tau spreading caused by cerebral amyloid accumulation.

## Declarations

### Ethics approval and consent to participate

N/A

## Consent for publication

N/A

## Availability of data and materials

All data generated or analyzed during this study are included in this published article [and its supplementary information files].

## Authors’ contributions

KRS, YX, CLC, and RV conceived and designed the study. YX designed and generated AAV. KRS, KPG, SC, MLL, AWK, JG performed experiments. KRS and KPG analyzed and interpreted data. KRS drafted manuscript. XY, CLC and RV edited manuscript. KRS, YX, CLC, and RV obtained funding. All authors read and approved submitted manuscript.

## Acknowledgements

Imaging work was performed at the Northwestern University Center for Advanced Microscopy (RRID: SCR_020996) generously supported by NCI CCSG P30 CA060553 awarded to the Robert H Lurie Comprehensive Cancer Center. mRNA sequencing was performed at the Northwestern University NUSeq Core Facility.

## Legends

**Supp Figure 1: High neuronal Nrf2 overexpression increases amyloid plaque load, neuroinflammation, and neuron loss in 5XFAD mice.** (**A**) Representative widefield microscopy images immunostained with NeuN (blue, neurons), Iba1 (green, microglia), GFAP (red, astrocytes) and MethoxyXO4 (white, dense core plaques). Size bar = 500μm. NB: Images are false colored for viewing. Imaging channels were as follows: MeXO4- 405nm, NeuN - 488nm, GFAP – 568nm, Iba1 – 647nm. Percent area covered by MeXO4 positive plaques (**B**) GFAP astrocytes (**C**) and Iba1 microglia (**D**) is significantly elevated in 5XFAD Nrf2 high transduced mice compared to untransduced in both cortex and hippocampus, while neuronal NeuN positive area (**E**) is decreased in 5XFAD Nrf2 high transduced mice in cortex and hippocampus, and in non-transgenic Nrf2 transduced mice in cortex. 2-way ANOVA with Sidak’s multiple comparisons test. * p<0.05, ** p<0.01, ***p<0.001, **** p<0.0001, one way ANOVA. Uninjected non-Tg *n=*6 (3F,3M), 5XFAD *n=*8 (4F,4M), Nrf2 high+GFP non-Tg *n=* 6 (3F, 3M), 5XFAD *n=*10 (5F, 5M).

## References

1. Aschenbrenner AJ, Gordon BA, Benzinger TLS, Morris JC, Hassenstab JJ. Influence of tau PET, amyloid PET, and hippocampal volume on cognition in Alzheimer disease. Neurology. Aug 28 2018;91(9):e859–e866. doi:10.1212/WNL.0000000000006075

2. Cho H, Choi JY, Lee HS, et al. Progressive Tau Accumulation in Alzheimer Disease: 2-Year Follow-up Study. J Nucl Med. Nov 2019;60(11):1611–1621. doi:10.2967/jnumed.118.221697

3. Hanseeuw BJ, Betensky RA, Jacobs HIL, et al. Association of Amyloid and Tau With Cognition in Preclinical Alzheimer Disease: A Longitudinal Study. JAMA Neurol. Aug 1 2019;76(8):915–924. doi:10.1001/jamaneurol.2019.1424

4. Hansson O. Biomarkers for neurodegenerative diseases. Nat Med. Jun 2021;27(6):954–963. doi:10.1038/s41591-021-01382-x

5. Jack CR, Jr., Knopman DS, Jagust WJ, et al. Hypothetical model of dynamic biomarkers of the Alzheimer’s pathological cascade. Lancet Neurol. Jan 2010;9(1):119–28. doi:10.1016/S1474-4422(09)70299-6

6. Jack CR, Jr., Wiste HJ, Schwarz CG, et al. Longitudinal tau PET in ageing and Alzheimer’s disease. Brain. May 1 2018;141(5):1517–1528. doi:10.1093/brain/awy059

7. Jagust W. Imaging the evolution and pathophysiology of Alzheimer disease. Nat Rev Neurosci. Nov 2018;19(11):687–700. doi:10.1038/s41583-018-0067-3

8. La Joie R, Visani AV, Baker SL, et al. Prospective longitudinal atrophy in Alzheimer’s disease correlates with the intensity and topography of baseline tau-PET. Sci Transl Med. Jan 1 2020;12(524)doi:10.1126/scitranslmed.aau5732

9. Mattsson-Carlgren N, Andersson E, Janelidze S, et al. Abeta deposition is associated with increases in soluble and phosphorylated tau that precede a positive Tau PET in Alzheimer’s disease. Sci Adv. Apr 2020;6(16):eaaz2387. doi:10.1126/sciadv.aaz2387

10. Ossenkoppele R, Rabinovici GD, Smith R, et al. Discriminative Accuracy of [18F]flortaucipir Positron Emission Tomography for Alzheimer Disease vs Other Neurodegenerative Disorders. JAMA. Sep 18 2018;320(11):1151–1162. doi:10.1001/jama.2018.12917

11. Palmqvist S, Insel PS, Stomrud E, et al. Cerebrospinal fluid and plasma biomarker trajectories with increasing amyloid deposition in Alzheimer’s disease. EMBO Mol Med. Dec 2019;11(12):e11170. doi:10.15252/emmm.201911170

12. Chen G, McKay NS, Gordon BA, et al. Predicting cognitive decline: Which is more useful, baseline amyloid levels or longitudinal change? Neuroimage Clin. 2024;41:103551. doi:10.1016/j.nicl.2023.103551

13. Mormino EC, Papp KV. Amyloid Accumulation and Cognitive Decline in Clinically Normal Older Individuals: Implications for Aging and Early Alzheimer’s Disease. J Alzheimers Dis. 2018;64(s1):S633–S646. doi:10.3233/JAD-179928

14. Karran E, De Strooper B. The amyloid hypothesis in Alzheimer disease: new insights from new therapeutics. Nat Rev Drug Discov. Apr 2022;21(4):306–318. doi:10.1038/s41573-022-00391-w

15. Boche D, Nicoll JAR. Invited Review - Understanding cause and effect in Alzheimer’s pathophysiology: Implications for clinical trials. Neuropathol Appl Neurobiol. Dec 2020;46(7):623–640. doi:10.1111/nan.12642

16. He Z, Guo JL, McBride JD, et al. Amyloid-beta plaques enhance Alzheimer’s brain tau-seeded pathologies by facilitating neuritic plaque tau aggregation. Nat Med. Jan 2018;24(1):29–38. doi:10.1038/nm.4443

17. Li T, Braunstein KE, Zhang J, et al. The neuritic plaque facilitates pathological conversion of tau in an Alzheimer’s disease mouse model. Nat Commun. Jul 4 2016;7:12082. doi:10.1038/ncomms12082

18. Alexander GC, Emerson S, Kesselheim AS. Evaluation of Aducanumab for Alzheimer Disease: Scientific Evidence and Regulatory Review Involving Efficacy, Safety, and Futility. JAMA. May 4 2021;325(17):1717–1718. doi:10.1001/jama.2021.3854

19. Mullard A. FDA approves third anti-amyloid antibody for Alzheimer disease. Nat Rev Drug Discov. Aug 2024;23(8):571. doi:10.1038/d41573-024-00116-1

20. Liu J, van Beusekom H, Bu XL, et al. Preserving cognitive function in patients with Alzheimer’s disease: The Alzheimer’s disease neuroprotection research initiative (ADNRI). Neuroprotection. Dec 2023;1(2):84–98. doi:10.1002/nep3.23

21. Shanks HRC, Chen K, Reiman EM, et al. p75 neurotrophin receptor modulation in mild to moderate Alzheimer disease: a randomized, placebo-controlled phase 2a trial. Nat Med. Jun 2024;30(6):1761–1770. doi:10.1038/s41591-024-02977-w

22. Nagahara AH, Mateling M, Kovacs I, et al. Early BDNF treatment ameliorates cell loss in the entorhinal cortex of APP transgenic mice. J Neurosci. Sep 25 2013;33(39):15596–602. doi:10.1523/JNEUROSCI.5195-12.2013

23. Tuszynski MH, Yang JH, Barba D, et al. Nerve Growth Factor Gene Therapy: Activation of Neuronal Responses in Alzheimer Disease. JAMA Neurol. Oct 2015;72(10):1139–47. doi:10.1001/jamaneurol.2015.1807

24. Brandes MS, Gray NE. NRF2 as a Therapeutic Target in Neurodegenerative Diseases. ASN Neuro. Jan-Dec 2020;12:1759091419899782. doi:10.1177/1759091419899782

25. Itoh K, Chiba T, Takahashi S, et al. An Nrf2/small Maf heterodimer mediates the induction of phase II detoxifying enzyme genes through antioxidant response elements. Biochem Biophys Res Commun. Jul 18 1997;236(2):313–22. doi:10.1006/bbrc.1997.6943

26. Itoh K, Wakabayashi N, Katoh Y, et al. Keap1 represses nuclear activation of antioxidant responsive elements by Nrf2 through binding to the amino-terminal Neh2 domain. Genes Dev. Jan 1 1999;13(1):76–86. doi:10.1101/gad.13.1.76

27. Motohashi H, Yamamoto M. Nrf2-Keap1 defines a physiologically important stress response mechanism. Trends Mol Med. Nov 2004;10(11):549–57. doi:10.1016/j.molmed.2004.09.003

28. Kensler TW, Wakabayashi N, Biswal S. Cell survival responses to environmental stresses via the Keap1-Nrf2-ARE pathway. Annu Rev Pharmacol Toxicol. 2007;47:89–116. doi:10.1146/annurev.pharmtox.46.120604.141046

29. Chorley BN, Campbell MR, Wang X, et al. Identification of novel NRF2-regulated genes by ChIP-Seq: influence on retinoid X receptor alpha. Nucleic Acids Res. Aug 2012;40(15):7416–29. doi:10.1093/nar/gks409

30. Liu T, Lv YF, Zhao JL, You QD, Jiang ZY. Regulation of Nrf2 by phosphorylation: Consequences for biological function and therapeutic implications. Free Radic Biol Med. May 20 2021;168:129–141. doi:10.1016/j.freeradbiomed.2021.03.034

31. Dhapola R, Beura SK, Sharma P, Singh SK, HariKrishnaReddy D. Oxidative stress in Alzheimer’s disease: current knowledge of signaling pathways and therapeutics. Mol Biol Rep. Jan 2 2024;51(1):48. doi:10.1007/s11033-023-09021-z

32. Perluigi M, Di Domenico F, Butterfield DA. Oxidative damage in neurodegeneration: roles in the pathogenesis and progression of Alzheimer disease. Physiol Rev. Jan 1 2024;104(1):103–197. doi:10.1152/physrev.00030.2022

33. Bahn G, Park JS, Yun UJ, et al. NRF2/ARE pathway negatively regulates BACE1 expression and ameliorates cognitive deficits in mouse Alzheimer’s models. Proc Natl Acad Sci U S A. Jun 18 2019;116(25):12516–12523. doi:10.1073/pnas.1819541116

34. Ramsey CP, Glass CA, Montgomery MB, et al. Expression of Nrf2 in neurodegenerative diseases. J Neuropathol Exp Neurol. Jan 2007;66(1):75–85. doi:10.1097/nen.0b013e31802d6da9

35. Jiwaji Z, Tiwari SS, Aviles-Reyes RX, et al. Reactive astrocytes acquire neuroprotective as well as deleterious signatures in response to Tau and Ass pathology. Nat Commun. Jan 10 2022;13(1):135. doi:10.1038/s41467-021-27702-w

36. Uruno A, Matsumaru D, Ryoke R, et al. Nrf2 Suppresses Oxidative Stress and Inflammation in App Knock-In Alzheimer’s Disease Model Mice. Mol Cell Biol. Feb 27 2020;40(6)doi:10.1128/MCB.00467-19

37. Yao Y, Ren Z, Yang R, et al. Salidroside reduces neuropathology in Alzheimer’s disease models by targeting NRF2/SIRT3 pathway. Cell Biosci. Nov 4 2022;12(1):180. doi:10.1186/s13578-022-00918-z

38. Branca C, Ferreira E, Nguyen TV, Doyle K, Caccamo A, Oddo S. Genetic reduction of Nrf2 exacerbates cognitive deficits in a mouse model of Alzheimer’s disease. Hum Mol Genet. Dec 15 2017;26(24):4823–4835. doi:10.1093/hmg/ddx361

39. Jo C, Gundemir S, Pritchard S, Jin YN, Rahman I, Johnson GV. Nrf2 reduces levels of phosphorylated tau protein by inducing autophagy adaptor protein NDP52. Nat Commun. Mar 25 2014;5:3496. doi:10.1038/ncomms4496

40. Calkins MJ, Vargas MR, Johnson DA, Johnson JA. Astrocyte-specific overexpression of Nrf2 protects striatal neurons from mitochondrial complex II inhibition. Toxicol Sci. Jun 2010;115(2):557–68. doi:10.1093/toxsci/kfq072

41. Gan L, Vargas MR, Johnson DA, Johnson JA. Astrocyte-specific overexpression of Nrf2 delays motor pathology and synuclein aggregation throughout the CNS in the alpha-synuclein mutant (A53T) mouse model. J Neurosci. Dec 5 2012;32(49):17775–87. doi:10.1523/JNEUROSCI.3049-12.2012

42. LaPash Daniels CM, Austin EV, Rockney DE, et al. Beneficial effects of Nrf2 overexpression in a mouse model of Alexander disease. J Neurosci. Aug 1 2012;32(31):10507–15. doi:10.1523/JNEUROSCI.1494-12.2012

43. Sigfridsson E, Marangoni M, Johnson JA, Hardingham GE, Fowler JH, Horsburgh K. Astrocyte-specific overexpression of Nrf2 protects against optic tract damage and behavioural alterations in a mouse model of cerebral hypoperfusion. Sci Rep. Aug 22 2018;8(1):12552. doi:10.1038/s41598-018-30675-4

44. Vargas MR, Johnson DA, Sirkis DW, Messing A, Johnson JA. Nrf2 activation in astrocytes protects against neurodegeneration in mouse models of familial amyotrophic lateral sclerosis. J Neurosci. Dec 10 2008;28(50):13574–81. doi:10.1523/JNEUROSCI.4099-08.2008

45. Cuadrado A, Rojo AI, Wells G, et al. Therapeutic targeting of the NRF2 and KEAP1 partnership in chronic diseases. Nat Rev Drug Discov. Apr 2019;18(4):295–317. doi:10.1038/s41573-018-0008-x

46. Xiong W, MacColl Garfinkel AE, Li Y, Benowitz LI, Cepko CL. NRF2 promotes neuronal survival in neurodegeneration and acute nerve damage. J Clin Invest. Apr 2015;125(4):1433–45. doi:10.1172/JCI79735

47. Wu DM, Ji X, Ivanchenko MV, et al. Nrf2 overexpression rescues the RPE in mouse models of retinitis pigmentosa. JCI Insight. Jan 25 2021;6(2)doi:10.1172/jci.insight.145029

48. Liang KJ, Woodard KT, Weaver MA, Gaylor JP, Weiss ER, Samulski RJ. AAV-Nrf2 Promotes Protection and Recovery in Animal Models of Oxidative Stress. Mol Ther. Mar 1 2017;25(3):765–779. doi:10.1016/j.ymthe.2016.12.016

49. Kanninen K, Heikkinen R, Malm T, et al. Intrahippocampal injection of a lentiviral vector expressing Nrf2 improves spatial learning in a mouse model of Alzheimer’s disease. Proc Natl Acad Sci U S A. Sep 22 2009;106(38):16505–10. doi:10.1073/pnas.0908397106

50. Karkkainen V, Pomeshchik Y, Savchenko E, et al. Nrf2 regulates neurogenesis and protects neural progenitor cells against Abeta toxicity. Stem Cells. Jul 2014;32(7):1904–16. doi:10.1002/stem.1666

51. Satoh T, Harada N, Hosoya T, Tohyama K, Yamamoto M, Itoh K. Keap1/Nrf2 system regulates neuronal survival as revealed through study of keap1 gene-knockout mice. Biochem Biophys Res Commun. Mar 6 2009;380(2):298–302. doi:10.1016/j.bbrc.2009.01.063

52. Salta E, Lazarov O, Fitzsimons CP, Tanzi R, Lucassen PJ, Choi SH. Adult hippocampal neurogenesis in Alzheimer’s disease: A roadmap to clinical relevance. Cell Stem Cell. Feb 2 2023;30(2):120–136. doi:10.1016/j.stem.2023.01.002

53. Terreros-Roncal J, Flor-Garcia M, Moreno-Jimenez EP, et al. Methods to study adult hippocampal neurogenesis in humans and across the phylogeny. Hippocampus. Apr 2023;33(4):271–306. doi:10.1002/hipo.23474

54. Yuan P, Zhang M, Tong L, et al. PLD3 affects axonal spheroids and network defects in Alzheimer’s disease. Nature. Dec 2022;612(7939):328–337. doi:10.1038/s41586-022-05491-6

55. Oakley H, Cole SL, Logan S, et al. Intraneuronal beta-amyloid aggregates, neurodegeneration, and neuron loss in transgenic mice with five familial Alzheimer’s disease mutations: potential factors in amyloid plaque formation. J Neurosci. Oct 4 2006;26(40):10129–40. doi:10.1523/JNEUROSCI.1202-06.2006

56. Gibson DG, Young L, Chuang RY, Venter JC, Hutchison CA, 3rd, Smith HO. Enzymatic assembly of DNA molecules up to several hundred kilobases. Nat Methods. May 2009;6(5):343–5. doi:10.1038/nmeth.1318

57. Grieger JC, Choi VW, Samulski RJ. Production and characterization of adeno-associated viral vectors. Nat Protoc. 2006;1(3):1412–28. doi:10.1038/nprot.2006.207

58. Xue Y, Wang SK, Rana P, et al. AAV-Txnip prolongs cone survival and vision in mouse models of retinitis pigmentosa. Elife. Apr 13 2021;10doi:10.7554/eLife.66240

59. Levites Y, Jansen K, Smithson LA, et al. Intracranial adeno-associated virus-mediated delivery of anti-pan amyloid beta, amyloid beta40, and amyloid beta42 single-chain variable fragments attenuates plaque pathology in amyloid precursor protein mice. J Neurosci. Nov 15 2006;26(46):11923–8. doi:10.1523/JNEUROSCI.2795-06.2006

60. Sadleir KR, Eimer WA, Kaufman RJ, Osten P, Vassar R. Genetic inhibition of phosphorylation of the translation initiation factor eIF2alpha does not block Abeta-dependent elevation of BACE1 and APP levels or reduce amyloid pathology in a mouse model of Alzheimer’s disease. PLoS One. 2014;9(7):e101643. doi:10.1371/journal.pone.0101643

61. Sadleir KR, Gomez KP, Edwards AE, et al. Annexin A6 membrane repair protein protects against amyloid-induced dystrophic neurites and tau phosphorylation in Alzheimer’s disease model mice. Acta Neuropathol. May 24 2025;149(1):51. doi:10.1007/s00401-025-02888-1

62. Martin M. Cutadapt removes adapter sequences from high-throughput sequencing reads. EMBnet J. 2011;17(1)

63. Putri GH, Anders S, Pyl PT, Pimanda JE, Zanini F. Analysing high-throughput sequencing data in Python with HTSeq 2.0. Bioinformatics. May 13 2022;38(10):2943–2945. doi:10.1093/bioinformatics/btac166

64. Mabrouk R, Miettinen PO, Tanila H. Most dystrophic neurites in the common 5xFAD Alzheimer mouse model originate from axon terminals. Neurobiol Dis. Jun 15 2023;182:106150. doi:10.1016/j.nbd.2023.106150

65. Sadleir KR, Kandalepas PC, Buggia-Prevot V, Nicholson DA, Thinakaran G, Vassar R. Presynaptic dystrophic neurites surrounding amyloid plaques are sites of microtubule disruption, BACE1 elevation, and increased Abeta generation in Alzheimer’s disease. Acta Neuropathol. Aug 2016;132(2):235–256. doi:10.1007/s00401-016-1558-9

66. Su JH, Cummings BJ, Cotman CW. Identification and distribution of axonal dystrophic neurites in Alzheimer’s disease. Brain Res. Oct 22 1993;625(2):228–37. doi:10.1016/0006-8993(93)91063-x

67. Tsering W, Prokop S. Neuritic Plaques - Gateways to Understanding Alzheimer’s Disease. Mol Neurobiol. May 2024;61(5):2808–2821. doi:10.1007/s12035-023-03736-7

68. Cras P, Kawai M, Lowery D, Gonzalez-DeWhitt P, Greenberg B, Perry G. Senile plaque neurites in Alzheimer disease accumulate amyloid precursor protein. Proc Natl Acad Sci U S A. Sep 1 1991;88(17):7552–6. doi:10.1073/pnas.88.17.7552

69. Cummings BJ, Su JH, Geddes JW, et al. Aggregation of the amyloid precursor protein within degenerating neurons and dystrophic neurites in Alzheimer’s disease. Neuroscience. Jun 1992;48(4):763–77. doi:10.1016/0306-4522(92)90265-4

70. Jorda-Siquier T, Petrel M, Kouskoff V, et al. APP accumulates with presynaptic proteins around amyloid plaques: A role for presynaptic mechanisms in Alzheimer’s disease? Alzheimers Dement. Nov 2022;18(11):2099–2116. doi:10.1002/alz.12546

71. Kandalepas PC, Sadleir KR, Eimer WA, Zhao J, Nicholson DA, Vassar R. The Alzheimer’s beta-secretase BACE1 localizes to normal presynaptic terminals and to dystrophic presynaptic terminals surrounding amyloid plaques. Acta Neuropathol. Sep 2013;126(3):329–52. doi:10.1007/s00401-013-1152-3

72. Zhao J, Fu Y, Yasvoina M, et al. Beta-site amyloid precursor protein cleaving enzyme 1 levels become elevated in neurons around amyloid plaques: implications for Alzheimer’s disease pathogenesis. J Neurosci. Apr 4 2007;27(14):3639–49.

73. Clark C, Lewczuk P, Kornhuber J, et al. Plasma neurofilament light and phosphorylated tau 181 as biomarkers of Alzheimer’s disease pathology and clinical disease progression. Alzheimers Res Ther. Mar 25 2021;13(1):65. doi:10.1186/s13195-021-00805-8

74. Suarez-Calvet M, Karikari TK, Ashton NJ, et al. Novel tau biomarkers phosphorylated at T181, T217 or T231 rise in the initial stages of the preclinical Alzheimer’s continuum when only subtle changes in Abeta pathology are detected. EMBO Mol Med. Dec 7 2020;12(12):e12921. doi:10.15252/emmm.202012921

75. Condello C, Schain A, Grutzendler J. Multicolor time-stamp reveals the dynamics and toxicity of amyloid deposition. Sci Rep. 2011;1:19. doi:10.1038/srep00019

76. Gowrishankar S, Yuan P, Wu Y, et al. Massive accumulation of luminal protease-deficient axonal lysosomes at Alzheimer’s disease amyloid plaques. Proc Natl Acad Sci U S A. Jul 14 2015;112(28):E3699–708. doi:10.1073/pnas.1510329112

77. Sanchez-Varo R, Trujillo-Estrada L, Sanchez-Mejias E, et al. Abnormal accumulation of autophagic vesicles correlates with axonal and synaptic pathology in young Alzheimer’s mice hippocampus. Acta Neuropathol. Jan 2012;123(1):53–70. doi:10.1007/s00401-011-0896-x

78. Nixon RA, Wegiel J, Kumar A, et al. Extensive involvement of autophagy in Alzheimer disease: an immuno-electron microscopy study. J Neuropathol Exp Neurol. Feb 2005;64(2):113–22. doi:10.1093/jnen/64.2.113

79. Fernandez-Valenzuela JJ, Sanchez-Varo R, Munoz-Castro C, et al. Enhancing microtubule stabilization rescues cognitive deficits and ameliorates pathological phenotype in an amyloidogenic Alzheimer’s disease model. Sci Rep. Sep 8 2020;10(1):14776. doi:10.1038/s41598-020-71767-4

80. Migheli A, Attanasio A, Pezzulo T, Gullotta F, Giordana MT, Schiffer D. Age-related ubiquitin deposits in dystrophic neurites: an immunoelectron microscopic study. Neuropathol Appl Neurobiol. Feb 1992;18(1):3–11. doi:10.1111/j.1365-2990.1992.tb00760.x

81. Torres M, Jimenez S, Sanchez-Varo R, et al. Defective lysosomal proteolysis and axonal transport are early pathogenic events that worsen with age leading to increased APP metabolism and synaptic Abeta in transgenic APP/PS1 hippocampus. Mol Neurodegener. Nov 22 2012;7:59. doi:10.1186/1750-1326-7-59

82. Caccamo A, Majumder S, Richardson A, Strong R, Oddo S. Molecular interplay between mammalian target of rapamycin (mTOR), amyloid-beta, and Tau: effects on cognitive impairments. J Biol Chem. Apr 23 2010;285(17):13107–20. doi:10.1074/jbc.M110.100420

83. Yang DS, Stavrides P, Mohan PS, et al. Reversal of autophagy dysfunction in the TgCRND8 mouse model of Alzheimer’s disease ameliorates amyloid pathologies and memory deficits. Brain. Jan 2011;134(Pt 1):258–77. doi:10.1093/brain/awq341

84. Jain A, Lamark T, Sjottem E, et al. p62/SQSTM1 is a target gene for transcription factor NRF2 and creates a positive feedback loop by inducing antioxidant response element-driven gene transcription. J Biol Chem. Jul 16 2010;285(29):22576–91. doi:10.1074/jbc.M110.118976

85. Forner S, Kawauchi S, Balderrama-Gutierrez G, et al. Systematic phenotyping and characterization of the 5xFAD mouse model of Alzheimer’s disease. Sci Data. Oct 15 2021;8(1):270. doi:10.1038/s41597-021-01054-y

86. Keren-Shaul H, Spinrad A, Weiner A, et al. A Unique Microglia Type Associated with Restricting Development of Alzheimer’s Disease. Cell. Jun 15 2017;169(7):1276–1290 e17. doi:10.1016/j.cell.2017.05.018

87. Krasemann S, Madore C, Cialic R, et al. The TREM2-APOE Pathway Drives the Transcriptional Phenotype of Dysfunctional Microglia in Neurodegenerative Diseases. Immunity. Sep 19 2017;47(3):566–581 e9. doi:10.1016/j.immuni.2017.08.008

88. Matsunaga E, Nambu S, Oka M, Tanaka M, Taoka M, Iriki A. Periostin, a neurite outgrowth-promoting factor, is expressed at high levels in the primate cerebral cortex. Dev Growth Differ. Apr 2015;57(3):200–8. doi:10.1111/dgd.12194

89. Tao Y, Dai P, Liu Y, et al. Erbin regulates NRG1 signaling and myelination. Proc Natl Acad Sci U S A. Jun 9 2009;106(23):9477–82. doi:10.1073/pnas.0901844106

90. Al-Saif A, Bohlega S, Al-Mohanna F. Loss of ERLIN2 function leads to juvenile primary lateral sclerosis. Ann Neurol. Oct 2012;72(4):510–6. doi:10.1002/ana.23641

91. Huang J, Tabbi-Anneni I, Gunda V, Wang L. Transcription factor Nrf2 regulates SHP and lipogenic gene expression in hepatic lipid metabolism. Am J Physiol Gastrointest Liver Physiol. Dec 2010;299(6):G1211–21. doi:10.1152/ajpgi.00322.2010

92. Liu S, Li X, Fan P, et al. The potential role of transcription factor sterol regulatory element binding proteins (SREBPs) in Alzheimer’s disease. Biomed Pharmacother. Nov 2024;180:117575. doi:10.1016/j.biopha.2024.117575

93. Ziegler AB, Thiele C, Tenedini F, et al. Cell-Autonomous Control of Neuronal Dendrite Expansion via the Fatty Acid Synthesis Regulator SREBP. Cell Rep. Dec 19 2017;21(12):3346–3353. doi:10.1016/j.celrep.2017.11.069

94. Spell C, Kolsch H, Lutjohann D, et al. SREBP-1a polymorphism influences the risk of Alzheimer’s disease in carriers of the ApoE4 allele. Dement Geriatr Cogn Disord. 2004;18(3-4):245–9. doi:10.1159/000080023

95. Bell KF, Al-Mubarak B, Martel MA, et al. Neuronal development is promoted by weakened intrinsic antioxidant defences due to epigenetic repression of Nrf2. Nat Commun. May 13 2015;6:7066. doi:10.1038/ncomms8066

96. Levings DC, Pathak SS, Yang YM, Slattery M. Limited expression of Nrf2 in neurons across the central nervous system. Redox Biol. Sep 2023;65:102830. doi:10.1016/j.redox.2023.102830

97. Claes C, Danhash EP, Hasselmann J, et al. Plaque-associated human microglia accumulate lipid droplets in a chimeric model of Alzheimer’s disease. Mol Neurodegener. Jul 23 2021;16(1):50. doi:10.1186/s13024-021-00473-0

98. Marschallinger J, Iram T, Zardeneta M, et al. Lipid-droplet-accumulating microglia represent a dysfunctional and proinflammatory state in the aging brain. Nat Neurosci. Feb 2020;23(2):194–208. doi:10.1038/s41593-019-0566-1

99. Huang H, Sharoar MG, Pathoulas J, et al. Accumulation of neutral lipids in dystrophic neurites surrounding amyloid plaques in Alzheimer’s disease. Biochim Biophys Acta Mol Basis Dis. Apr 2024;1870(4):167086. doi:10.1016/j.bbadis.2024.167086

100. Yao Y, Nzou G, Alle T, et al. Correction of microtubule defects within Abeta plaque-associated dystrophic axons results in lowered Abeta release and plaque deposition. Alzheimers Dement. Oct 2020;16(10):1345–1357. doi:10.1002/alz.12144

101. Zhang B, Yao Y, Cornec AS, et al. A brain-penetrant triazolopyrimidine enhances microtubule-stability, reduces axonal dysfunction and decreases tau pathology in a mouse tauopathy model. Mol Neurodegener. Nov 7 2018;13(1):59. doi:10.1186/s13024-018-0291-3

102. Brunden KR, Lee VM, Smith AB, 3rd, Trojanowski JQ, Ballatore C. Altered microtubule dynamics in neurodegenerative disease: Therapeutic potential of microtubule-stabilizing drugs. Neurobiol Dis. Sep 2017;105:328–335. doi:10.1016/j.nbd.2016.12.021

103. Yao Y, Muench M, Alle T, et al. A small-molecule microtubule-stabilizing agent safely reduces Abeta plaque and tau pathology in transgenic mouse models of Alzheimer’s disease. Alzheimers Dement. Jul 2024;20(7):4540–4558. doi:10.1002/alz.13875

104. Gumy LF, Chew DJ, Tortosa E, et al. The kinesin-2 family member KIF3C regulates microtubule dynamics and is required for axon growth and regeneration. J Neurosci. Jul 10 2013;33(28):11329–45. doi:10.1523/JNEUROSCI.5221-12.2013

105. Guzik-Lendrum S, Rayment I, Gilbert SP. Homodimeric Kinesin-2 KIF3CC Promotes Microtubule Dynamics. Biophys J. Oct 17 2017;113(8):1845–1857. doi:10.1016/j.bpj.2017.09.015

106. Guillabert-Gourgues A, Jaspard-Vinassa B, Bats ML, et al. Kif26b controls endothelial cell polarity through the Dishevelled/Daam1-dependent planar cell polarity-signaling pathway. Mol Biol Cell. Mar 15 2016;27(6):941–53. doi:10.1091/mbc.E14-08-1332

107. Wojcik MH, Okada K, Prabhu SP, et al. De novo variant in KIF26B is associated with pontocerebellar hypoplasia with infantile spinal muscular atrophy. Am J Med Genet A. Dec 2018;176(12):2623–2629. doi:10.1002/ajmg.a.40493

108. Castanho I, Yeganeh PN, Boix CA, et al. Molecular hallmarks of excitatory and inhibitory neuronal resilience and resistance to Alzheimer’s disease. bioRxiv. Jan 15 2025;doi:10.1101/2025.01.13.632801

109. Haass C, Lemere CA, Capell A, et al. The Swedish mutation causes early-onset Alzheimer’s disease by beta-secretase cleavage within the secretory pathway. Nat Med. Dec 1995;1(12):1291–6. doi:10.1038/nm1295-1291

110. Sadleir KR, Eimer WA, Cole SL, Vassar R. Abeta reduction in BACE1 heterozygous null 5XFAD mice is associated with transgenic APP level. Mol Neurodegener. Jan 7 2015;10:1. doi:10.1186/1750-1326-10-1

111. Long Z, Chen J, Zhao Y, et al. Dynamic changes of autophagic flux induced by Abeta in the brain of postmortem Alzheimer’s disease patients, animal models and cell models. Aging (Albany NY). Jun 13 2020;12(11):10912–10930. doi:10.18632/aging.103305

112. He X, Dando O, Qiu J. Nrf2 controls homeostatic transcriptional signatures and inflammatory responses in a cell-type specific manner in the adult mouse brain. iScience. Sep 19 2025;28(9):113198. doi:10.1016/j.isci.2025.113198

113. Kao YC, Ho PC, Tu YK, Jou IM, Tsai KJ. Lipids and Alzheimer’s Disease. Int J Mol Sci. Feb 22 2020;21(4)doi:10.3390/ijms21041505

114. Zarrouk A, Debbabi M, Bezine M, et al. Lipid Biomarkers in Alzheimer’s Disease. Curr Alzheimer Res. Feb 22 2018;15(4):303–312. doi:10.2174/1567205014666170505101426

115. He S, Xu Z, Han X. Lipidome disruption in Alzheimer’s disease brain: detection, pathological mechanisms, and therapeutic implications. Mol Neurodegener. Jan 27 2025;20(1):11. doi:10.1186/s13024-025-00803-6

116. Ebner M, Frohlich F, Haucke V. Mechanisms and functions of lysosomal lipid homeostasis. Cell Chem Biol. Mar 20 2025;32(3):392–407. doi:10.1016/j.chembiol.2025.02.003

117. Mrdenovic D, Pieta IS, Nowakowski R, Kutner W, Lipkowski J, Pieta P. Amyloid beta interaction with model cell membranes - What are the toxicity-defining properties of amyloid beta? Int J Biol Macromol. Mar 1 2022;200:520–531. doi:10.1016/j.ijbiomac.2022.01.117

