## supplemental figure 1 for "Neuronal overexpression of Nrf2 reduces dystrophic neurites in 5XFAD Alzheimer’s disease model mice"

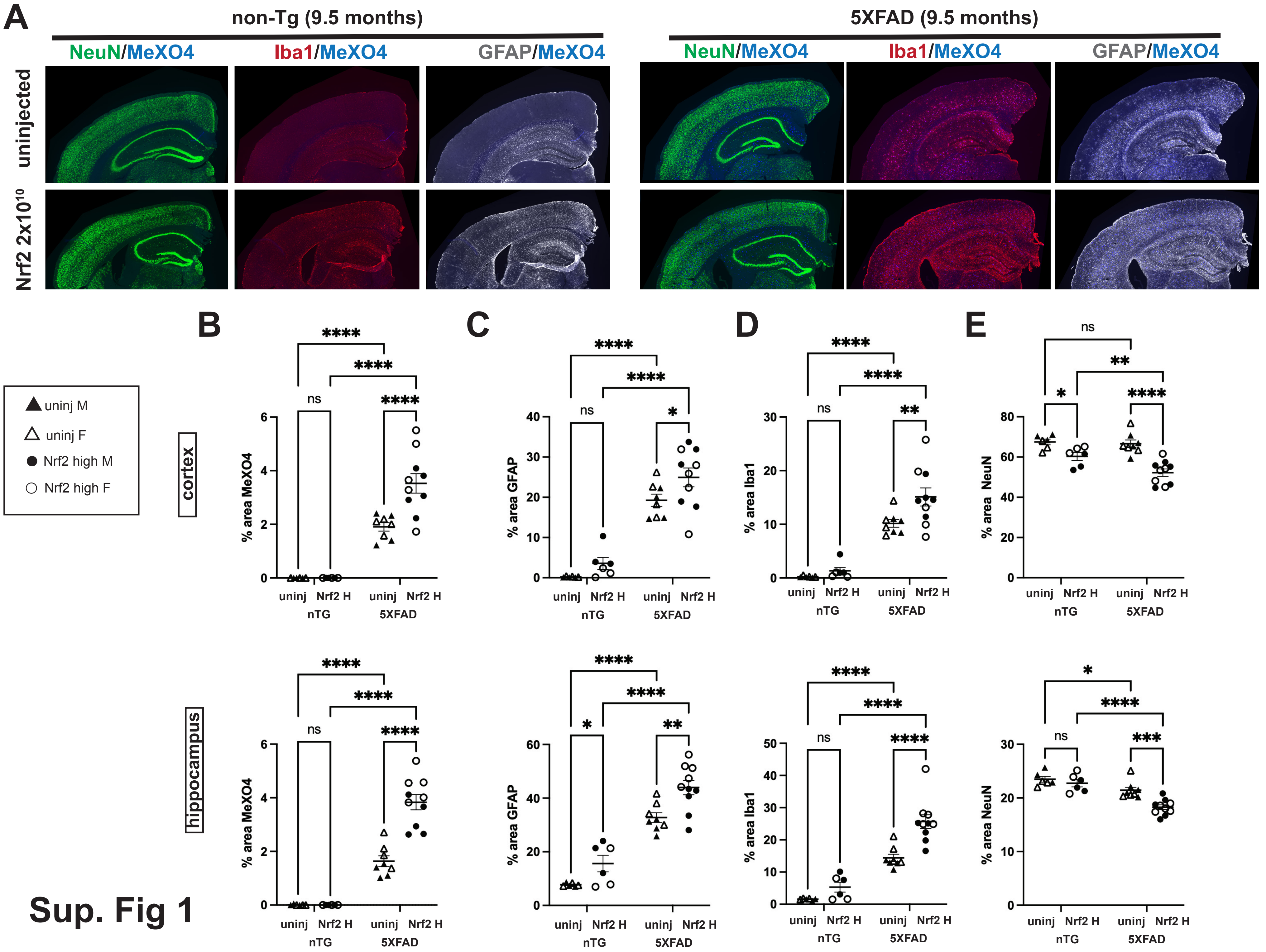
A
non-Tg (9.5 months)
5XFAD (9.5 months)
NeuN/MeXO4
Iba1/MeXO4
GFAP/MeXO4
uninjected
Nrf2 2x10^10
B
C
D
E
uninj M
uninj F
Nrf2 high M
Nrf2 high F
cortex
hippocampus
% area MeXO4
% area GFAP
% area Iba1
% area NeuN
uninj
Nrf2 H
nTG
5XFAD
ns
****
**
***
*

Sup. Fig 1

**Supp Figure 1: High neuronal Nrf2 overexpression increases amyloid plaque load, neuroinflammation, and neuron loss in 5XFAD mice.** (**A**) Representative widefield microscopy images immunostained with NeuN (blue, neurons), Iba1 (green, microglia), GFAP (red, astrocytes) and MethoxyXO4 (white, dense core plaques). Size bar = 500µm. NB: Images are false colored for viewing. Imaging channels were as follows: MeXO4- 405nm, NeuN - 488nm, GFAP – 568nm, Iba1 – 647nm. Percent area covered by MeXO4 positive plaques (**B**) GFAP astrocytes (**C**) and Iba1 microglia (**D**) is significantly elevated in 5XFAD Nrf2 high transduced mice compared to untransduced in both cortex and hippocampus, while neuronal NeuN positive area (**E**) is decreased in 5XFAD Nrf2 high transduced mice in cortex and hippocampus, and in non-transgenic Nrf2 transduced mice in cortex. 2-way ANOVA with Sidak's multiple comparisons test. * $p<0.05$, ** $p<0.01$, ***$p<0.001$, **** $p<0.0001$, one way ANOVA. Uninjected non-Tg *n*=6 (3F,3M), 5XFAD *n*=8 (4F,4M), Nrf2 high+GFP non-Tg *n*= 6 (3F, 3M), 5XFAD *n*=10 (5F, 5M).
